# Specialized neurons in the right habenula mediate response to aversive olfactory cues

**DOI:** 10.1101/2021.07.19.452986

**Authors:** Jung-Hwa Choi, Erik Duboué, Michelle Macurak, Jean-Michael Chanchu, Marnie E. Halpern

## Abstract

Hemispheric specializations are well studied at the functional level but less is known about the underlying neural mechanisms. We identified a small cluster of cholinergic neurons in the right dorsal habenula (dHb) of zebrafish, defined by their expression of the *lecithin retinol acyltransferase domain containing 2a* (*lratd2a*) gene and their efferent connections with a subregion of the ventral interpeduncular nucleus (vIPN). The unilateral *lratd2a*-expressing neurons are innervated by a subset of mitral cells from both the left and right olfactory bulb and are activated upon exposure of adult zebrafish to the aversive odorant cadaverine that provokes avoidance behavior. Using an intersectional strategy to drive expression of the botulinum neurotoxin specifically in these neurons, we find that adults no longer show protracted avoidance to cadaverine. Mutants with left-isomerized dHb that lack these neurons are less repelled by cadaverine and their behavioral response to alarm substance, a potent aversive cue, is diminished. However mutants in which both dHb have right identity appear more reactive to alarm substance. The results implicate an asymmetric dHb-vIPN neural circuit in processing of aversive olfactory cues and modulating resultant behavioral responses.

## Introduction

Fish use the sense of smell to search for food, detect danger, navigate and communicate social information by detecting chemical cues in their aquatic environment (Yoshihara, 2014). As with birds and mammals, perception of olfactory cues is lateralized and influences behavior (Siniscalchi, 2017). In zebrafish, nine glomerular clusters in the olfactory bulb (OB) receive olfactory information from sensory neurons in the olfactory epithelium and, in turn, transmit signals to four forebrain regions: the posterior zone of the dorsal telencephalon (Dp), the ventral nucleus of the ventral telencephalon (Vv), the posterior tuberculum (PT), and the dorsal habenular region (dHb) (Miyasaka et al., 2014; Yoshihara, 2014). In contrast to all other target regions that are located on both sides of the forebrain, only the right nucleus of the dHb is innervated by mitral cells that emanate from medio-dorsal (mdG) and ventro-medial (vmG) glomerular clusters in both OBs (Miyasaka *et al*., 2014; Yoshihara, 2014). Moreover, calcium imaging experiments suggest that the right dHb shows a preferential response to odorants compared to the left dHb (Chen et al., 2019; Dreosti et al., 2014; Jetti et al., 2014; Krishnan et al., 2014). The identity of the post-synaptic neurons within the right dHb that receive olfactory input and the purpose of this asymmetric connection are unknown.

The habenulae are highly conserved structures in the vertebrate brain and, in teleosts such as zebrafish, consist of dorsal and ventral (vHb) nuclei, which are equivalent to the medial and lateral habenulae of mammals, respectively (Amo et al., 2010). The neurons of the dHb are largely glutamatergic and contain specialized subpopulations that also produce acetylcholine, substance P or somatostatin. In zebrafish, the number of neurons within each subtype differs between the left and right dHb (deCarvalho et al., 2014; Hsu et al., 2016). The dHb have been implicated in diverse states such as reward, fear, anxiety, sleep and addiction (Duboué et al., 2017; Hikosaka, 2010; Okamoto et al., 2012). Accordingly, the right dHb was shown to respond to bile acid and involved in food-seeking behaviors (Chen *et al*., 2019; Krishnan *et al*., 2014), whereas the left dHb was found to be activated by light and attenuate fear responses (Dreosti *et al*., 2014; Facchin et al., 2015; Zhang et al., 2017). However, the properties of the dHb neurons implicated in these behaviors, such as their neurotransmitter identity and precise connectivity with their unpaired target, the midbrain interpeduncular nucleus (IPN), have yet to be determined.

Here, we describe a group of cholinergic neurons defined by their expression of the *lecithin retinol acyltransferase domain containing 2a* (*lratd2a*) gene [formerly known as *family with sequence similarity 84 member B* (*fam84b*)], that are located in the right dHb where they are selectively innervated by the olfactory mitral cells that originate from both sides of the brain (Miyasaka et al., 2009), and form efferent connections with a restricted subregion of the ventral IPN (vIPN). Activity of the *lratd2a*-expressing neurons is increased following exposure to the aversive odorant cadaverine and their inactivation alters the avoidance behavior of adult zebrafish to this repulsive cue. Our findings provide further evidence for functional specialization of the left and right habenular nuclei and reveal the neuronal pathway that mediates a lateralized olfactory response.

## Materials and Methods

### Zebrafish

Zebrafish were maintained at 27 °C in a 14:10 h light/dark cycle in a recirculating system with dechlorinated water (system water). The AB wild-type strain (Walker, 1998), transgenic lines *Tg(lratd2a:QF2)*^*c601*^, *Tg(slc5a7a:Cre)*^*c662*^, *Tg(Xla*.*Tubb2:QF2;he1*.*1:mCherry)*^*c663*^, *Tg(QUAS:GCaMP6f)*^*c587*^, *Tg(QUAS:BoTxBLC-GFP)*^*c605*^, *Tg(QUAS:mApple-CAAX;he1*.*1:mCherry)*^*c636*^, *Tg(QUAS:loxP-mCherry-loxP-GFP-CAAX)*^*c679*^, and *Tg(QUAS:loxP-mCherry-loxP-BoTxBLC-GFP)*^*c674*^, *Tg(−10lhx2a:gap-EYFP)*^*zf177*^ (formally known as *Tg(lhx2a:gap-YFP)*) (Miyasaka *et al*., 2009), and mutant strains *tcf7l2*^*zf55*^ (Muncan et al., 2007) and *bsx*^*m1376*^ (Schredelseker and Driever, 2018) were used. For imaging, embryos and larvae were transferred to system water containing 0.003% phenylthiourea (PTU) to inhibit melanin pigmentation. All zebrafish protocols were approved by the Institutional Animal Care and Use Committee (IACUC) of the Carnegie Institution for Science or Dartmouth College.

### Generation of transgenic lines by *Tol2* transgenesis

The MultiSite Gateway-based construction kit (Kwan et al., 2007) was used to create transgenic constructs for Tol2 transposition. A 16 bp *QUAS* sequence (Potter et al., 2010), was cloned into the 5’ entry vector (*pDONRP4-P1R*, #219 of Tol2kit v1.2) via a BP reaction (11789020, Thermo Fisher Scientific). Middle entry vectors (*pDONR221*, #218 of Tol2kit v1.2 (Kwan *et al*., 2007)) were generated for *QF2, mApple-CAAX, loxP-mCherry-stop-loxP, GCaMP6f* and *BoTxBLC-GFP*. Sequences corresponding to the *SV40 poly A* tail, the *SV40 poly A* tail followed by a secondary marker consisting of the zebrafish *hatching enzyme 1* promoter (Xie et al., 2012) driving mCherry, or to *BoTxBLC-GFP* (Lal et al., 2018; Sternberg et al., 2016; Zhang *et al*., 2017) were cloned into the 3’ entry vector (*pDONRP2R-P3*, #220 of Tol2kit v1.2 (Kwan *et al*., 2007)). Final constructs were created using an LR reaction (11791020, Thermo Fisher Scientific) into a Tol2 destination vector (*pDestTol2pA2*, #394 of the Tol2kit v1.2 (Kwan *et al*., 2007)) (Supplementary table 1).

To produce Tol2 transposase mRNA, *pCS-zT2TP* was digested by *Not*I and RNA synthesized using the mMESSAGE mMACHINE SP6 Transcription Kit (AM1340, Thermo Fisher Scientific). RNA was purified by phenol/chloroform-isoamyl extraction, followed by chloroform extraction and isopropanol precipitation (Suster et al., 2011). A solution containing *QF2/QUAS* plasmid DNA (∼25 ng/μl), transposase mRNA (∼25 ng/μl) and phenol red (0.5%) was microinjected into one-cell stage zebrafish embryos, which were raised to adulthood. To identify transgenic founders, F_0_ adult fish were outcrossed to AB and embryos were assessed for the presence of the secondary marker by screening for mCherry labeling of hatching gland cells after 24 hpf and raised to adulthood.

### Generation of transgenic lines by genome editing

For generating transgenic lines at targeted sites, we performed CRISPR/Cas9-mediated genome editing using the method of Kimura *et al*. (Kimura et al., 2014), which relies on homology-independent repair of double-strand breaks for integration of donor DNA. To construct the donor DNA, we combined GFP bait sequences (Gbait) and the hsp70 promoter fragment (Kimura *et al*., 2014), with a QF2 sequence, which contains the DNA binding and transcriptional activation domains of the QF transcription factor of *Neurospora crassa* (Ghosh and Halpern, 2016; Subedi et al., 2014). The Gbait-hsp70 sequence was amplified with forward 5’-GGCGAGGGCGATGCCACCTACGG-3’ and reverse 5’-CCGCGGCAAGAAACTGCAATAAAAAAAAC-3’ primers, using Gbait-hsp70:Gal4 donor DNA (Kimura *et al*., 2014). QF2 sequence was amplified with forward 5’-ACTAGTATGCCACCCAAGCGCAAAACGC-3’ and reverse 5’-CTGCAGCAACTATGTATAATAAAGTTGAAA-3’ primers, using pDEST:QF2 template DNA Subsequently, the Gbait-hsp70 fragment and QF2 fragment were independently inserted into pGEM T-easy (A1360, Promega) and subsequently combined into one vector by *Sac*II digestion and ligation (Addgene, plasmid #122563). The Cre sequence was amplified using pCR8GW-Cre-pA-FRT-kan-FRT as template DNA (Suster *et al*., 2011) and (forward 5’-ACTAGTGCCACCATGGCCAATTTACTG-3’, and reverse 5’-CTGCAGGGACAAACCACAACTAGA-3’) primers, and inserted into pGEM T-easy. The Gbait-hsp70 fragment was subcloned into the Cre vector by *Sac*II digestion and ligation (Addgene, plasmid #122562).

Production of sgRNAs and Cas9 RNA was performed as described previously (Hwang et al., 2013; Jao et al., 2013). Potential sgRNAs were designed using Zifit (Sander et al., 2010). Pairs of synthetic oligonucleotides (*lratd2a* sense, 5’-TAGGACTGGACACCGAAGAAGA-3’; *lratd2a* anti-sense, 5’-AAACTCTTCTTCGGTGTCCAGT-3’; *slc5a7a* sense, 5’-TAGGCTCTTTGTGCACTGTTGG-3’; *slc5a7a* anti-sense, 5’-AAACCCAACAGTGCACAAAGAG-3’), 5’-TAGG-N_18_-3’ and 5’-AAAC-N_18_-3’, were annealed and inserted at the *Bsa*I site of the pDR274 vector (Addgene, plasmid #42250). To make sgRNA and Cas9 mRNA, template DNA for sgRNAs and pT3TS nCas9n (Addgene, plasmid #46757) were digested by *Dra*I and *Xba*I, respectively. The MAXIscript T7 Transcription Kit (AM1312, Thermo Fisher Scientific) was used for synthesis of sgRNAs from linearized DNA template and the mMESSAGE mMACHINE T3 Transcription Kit (AM1348, Thermo Fisher Scientific) for synthesis of Cas9 RNA. RNA was purified by phenol/chloroform and precipitated by isopropanol.

A solution containing sgRNA for the targeted gene (∼50 ng/μl), sgRNA (∼50 ng/μl) to linearize donor plasmids at the Gbait site (Auer et al., 2014; Kimura *et al*., 2014), the Gbait-hsp70-QF2-pA and Gbait-hsp70-Cre-pA (∼50 ng/μl) plasmids, Cas9 mRNA (∼500 ng/μl), and phenol red (0.5%) was microinjected into one-cell stage embryos. To verify integration of donor DNA at the target locus, PCR was performed using primers that correspond to sequences flanking the integration site and within the donor plasmid (*hsp70* reverse, 5’-TCAAGTCGCTTCTCTTCGGT-3’). (For *lratd2a, the* forward primer is 5’-CTGCTGAAGTGGCATTTATGGGC-3’ and the reverse primer is 5’-CCTGGAAGTCCCCGACATAC-3’; for *slc5a7a* the forward primer is 5’-CACATCTCTCTGACGTCCATC-3’ and the reverse is 5’-GTTGCTGCGCAGGACTTAAAA-3’). Sequence analysis of PCR products confirmed integration at the targeted sites.

### RNA *in situ* hybridization

Whole-mount *in situ* hybridization was performed as previously described (deCarvalho *et al*., 2014; Gamse et al., 2002). In brief, larvae and dissected brains were fixed in 4% paraformaldehyde (P6148, Sigma-Aldrich) in 1X PBS (phosphate-buffered saline) at 4 °C overnight. To synthesize RNA probes, the following restriction enzymes and RNA polymerases were used: *lratd2a* (*Bam*HI/T7), *fos* (*Not*I/SP6), *slc5a7a* (*Not*I/SP6), *kctd12*.*1* (*Eco*RI/T7) (deCarvalho et al., 2013; Hong et al., 2013). Probes were labeled with UTP-digoxigenin (11093274910, Roche) and samples incubated at 70 °C in hybridization solution containing 50% formamide. Hybridized probes were detected using alkaline phosphatase-conjugated antibodies (Anti-Digoxigenin-AP, #11093274910, and Anti-Fluorescein-AP, #11426338910, Sigma-Aldrich) and visualized by staining with 4-nitro blue tetrazolium (NBT, #11383213001, Roche), 5-bromo-4-chloro-3-indolyl-phosphate (BCIP, #11383221001, Roche) and 2-(4-Iodophenyl)-3-(4-nitrophenyl)-5-phenyltetrazolium Chloride (INT, #I00671G, Fisher Scientific).

### Preparation of odorants

Alarm substance was freshly prepared on the day of testing. Adult zebrafish (6 female and 6 male) were anesthetized in 0.02% tricaine (E10521, Ethyl 3-aminobenzoate methanesulfonate; Sigma-Aldrich). Shallow lesions were made on the skin (10 on each side) using a fresh razor blade and single fish were consecutively immersed in a beaker containing distilled water (25 ml for *fos* experiment and 50 ml for behavioral analyses) for 30 seconds at 4 °C. The solution was filtered using a 0.2 μm filter (565-0020, ThermoFisher Scientific) and stored at 4 °C until used (Mathuru et al., 2012). The cadaverine (#33211, final concentration 100 μM) and chondroitin sulfate (#c4384, 100 μg/ml) were purchased from Sigma-Aldrich, and stock solutions prepared in distilled water.

### Calcium imaging in larval and juvenile zebrafish

Calcium imaging was performed on 7, 14 and 21-22 dpf *Tg(lratd2a:QF2)*^*c644*^; *Tg(QUAS:GCaMP6f)*^*c587*^ individuals. Larvae and juveniles were paralyzed by immersion in α-bungarotoxin (20 µl of 1 mg/ml solution in system water, B1601, ThermoFisher Scientific) (Duboué *et al*., 2017; Severi et al., 2014) followed by washing in fresh system water. Individual fish were embedded in 2% low melting agarose in a petri dish (60 mm) with a custom-designed mold. After solidification, the agarose around the nose was carefully removed with forceps for access to odorants, and the individual immersed in fresh system water. The dish was placed under a 25X (NA = 0.95), on a Leica SP5 (for chondroitin sulfate) or under a 20X (NA=0.5) water immersion objective on a Zeiss LSM 980 (for cadaverine) confocal microscope. Images were acquired in *XYZT* acquisition mode at 512 × 200 pixel resolution at a rate of 2 Hz and digitized 8 bit from two focal planes. To calculate fluorescence intensity, regions of interest (ROI) were manually drawn around each cell in the average focal plane with the polygon tool and ROI *manage* in Fiji (Schindelin et al., 2012). To normalize calcium activity for each cell to baseline fluorescence (average of 250 frames from each neuron), the fractional change in fluorescence (ΔF/F) was calculated before the application of odorants, according to the formula F = (F_i_-F_0_)/F_0_, where F_i_ is the fluorescence intensity at a single time point and F_0_ is the baseline fluorescence. All data and images were analyzed using custom programs in MATLAB (MathWorks, version 7.3) and Excel software.

### Assay of *fos* expression in adult zebrafish

Individual adult zebrafish (7–9 months old) were placed in a tank with 1 L system water and acclimated for at least 1 hour prior to odorant exposure. Each odorant solution (1 ml) was gently pipetted into the tank water and the fish was kept there as the odorant diffused. After 30 min, the fish was sacrificed in an ice water slurry, and the brain dissected out and fixed in 4% paraformaldehyde in 1X PBS overnight at 4 °C. Fixed brains were embedded in 4% low melting agarose (SeaPlaque® Agarose, Lonza) in 1X PBS and sectioned at 50 μm (for juvenile brains) or 70 μm (for adult brains) using a vibratome (VT1000S, Leica Biosystems, Inc.). For more precise counting of *fos* expressing cells in adult brains, habenular sections were 35 μm thick. Sections were covered in 50% glycerol in 1X PBS under coverslips. Bright-field images were captured with a Zeiss AxioCam HRc camera mounted on a Zeiss Axioskop. A Leica SP5 confocal microscope was used for fluorescent images. Data from *fos* RNA *in situ* hybridization experiments were quantified using ImageJ/Fiji software (Schindelin *et al*., 2012).

### Behavioral assays

Behavioral assays were performed using 5 – 7 week old juvenile zebrafish and adults between 4 and 8 months of age. Responses to odorants were measured between 10:00 a.m. and 4:00 p.m. and fish were starved for 1 day prior to testing (Koide et al., 2009). Individual adults were placed in a 1.5 L test tank (Aquatic Habitats) in 1 L of system water and allowed to acclimate for at least 1 hour. For experiments with juveniles, individuals were acclimated to the behavior room for 1 hour, gently netted into the test tank (20 × 9 × 8.3 cm, 1.5 L mating cage) with 0.6 L system water and maintained there for 5 min prior to testing. Swimming activity was recorded for 5 min (4 min for juveniles) before and after the application of odorants. Odorants (2 ml for adults, 1 ml for juveniles) were slowly expelled through plastic tubing (Tygon R-3606; 0.8 mm ID, 2.4 mm OD) attached on one end to a 3 ml syringe (BD 309657) and on the other positioned at one end of the test tank. Preference indices were calculated using the formula: preference to odorants = (Total time spent in the tank half where odorant was delivered) − (Total time spent in the other half of the tank)/Total time (Koide *et al*., 2009; Wakisaka et al., 2017).

### Quantification and statistical analyses

Analyses were performed using custom written scripts in MATLAB (The MathWorks). Mann Whitney *U* and Student’s *t* tests were used to compare nonparametric unmatched groups. One-sample *t* test against 0 was used for analyzing the preference to cadaverine. The significance was two-tailed for all tests and depicted as n.s. (non-significant, P> 0.05) or with significance as *P<0.05, **P<0.01, ***P<0.001, and ****P<0.0001.

## Results

### lratd2a-expressing neurons in the right dHb receive bilateral olfactory input

A subset of medio-dorsal mitral cells that are labeled by *Tg(lhx2a:gap-YFP)* were previously shown to project their axons bilaterally through the telencephalon and terminate in the right dHb (Miyasaka *et al*., 2009), in the vicinity of a small population of *lratd2a*-expressing neurons [(deCarvalho *et al*., 2013) and Figure 1A]. To characterize this subset of right dHb neurons, we used CRISPR/Cas9-mediated targeted integration (Kimura *et al*., 2014) to introduce the QF2 transcription factor (Ghosh and Halpern, 2016; Subedi *et al*., 2014) under the control of *lratd2a* regulatory sequences (Figure 1B). QF2 does not disrupt transcription at the *lratd2a* locus (Supplementary figure 1) and drives expression of QUAS regulated fluorescent reporter genes in a similar pattern as endogenous gene expression in the nervous system (deCarvalho *et al*., 2013). In 4 dpf larval zebrafish, this includes a subset of neurons in the OB, the bilateral vHb, and a small cluster of neurons in the right dHb (Figures 1C-E’’). Double labeling confirms that the axons of *lhx2a* olfactory neurons terminate precisely at the *lratd2a*-expressing right habenular neurons (Figures 1C-D’’), which project to a restricted region of the ventral IPN (Figures 1F-G’’).

**Figure 1.**
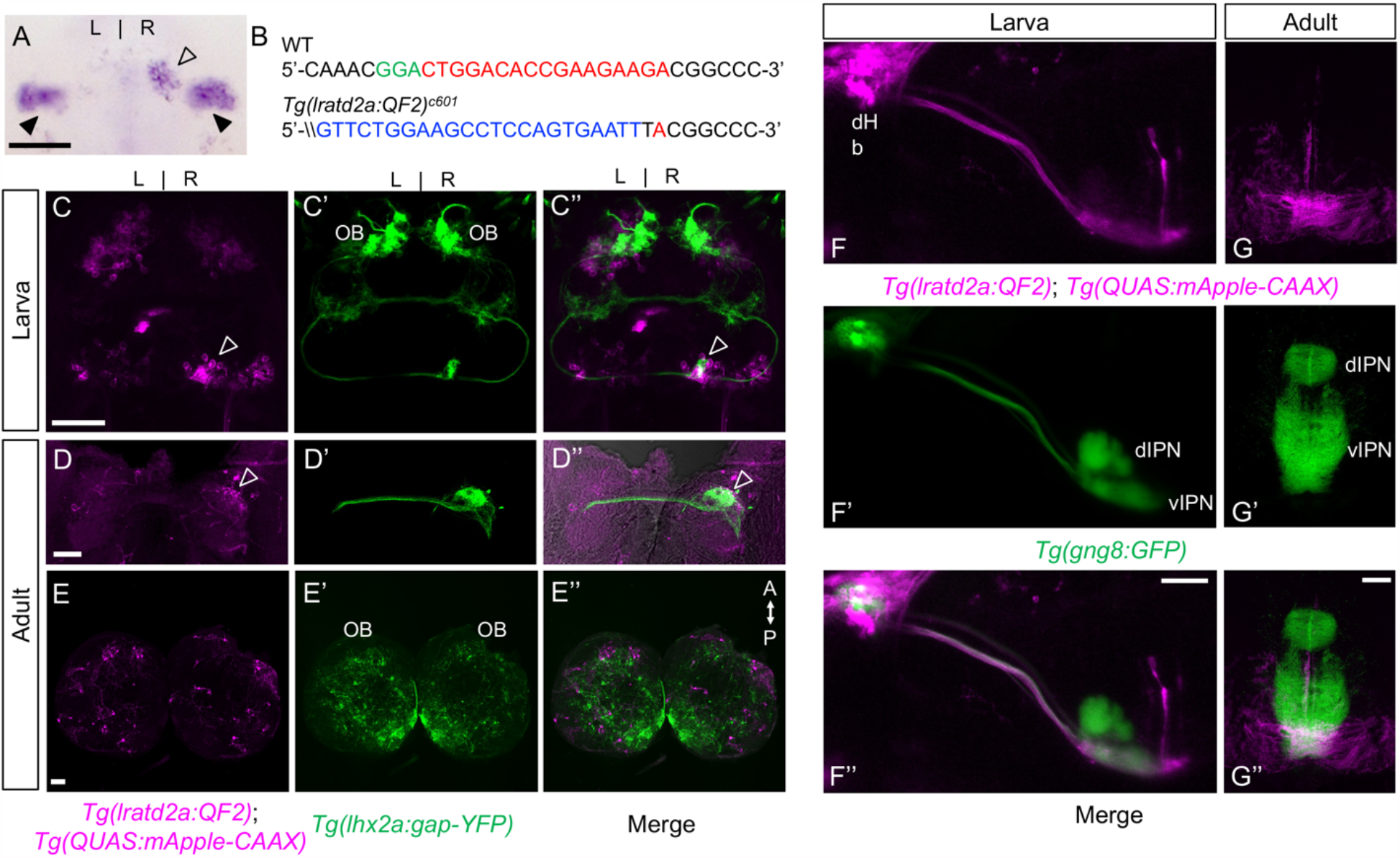
*lratd2a*-expressing neurons in the right dHb connect asymmetric pathway from the olfactory bulb to ventral IPN. (A) Pattern of *lratd2a* expression at 5 days post fertilization (dpf), open arrowhead indicates right dHb and black arrowheads the bilateral vHb. (B) Sequences of WT (top) and transgenic fish (bottom) with QF2 integrated within the first exon of the *lratd2a* gene. PAM sequences are green, the sgRNA binding site red and donor DNA blue. Confocal dorsal views of *Tg(lratd2a:QF2), Tg(QUAS:mApple-CAAX)* and *Tg(lhx2a:gap-YFP)* labeling in a (C-C’’) 5 dpf larva and in transverse sections of the adult brain at 3 months post-fertilization (mpf) at the level of the (D-D’’) dHb and (E-E’’) olfactory bulb. Axons of *lhx2a* olfactory mitral cells (open arrowheads, C and D) terminate at *lratd2a* dHb neurons. (F-F’’) Lateral view of *Tg(lratd2a:QF2), Tg(QUAS:mApple-CAAX), Tg(gng8:GFP)* larva at 6 dpf with mApple labeled dHb terminals at the ventral interpeduncular nucleus (vIPN). Dorsal habenular nuclei (dHb), dorsal interpeduncular nucleus (dIPN). (G-G’’) Axonal endings of *lratd2a* dHb neurons are restricted to the ventralmost region of the vIPN in transverse section of 2.5 mpf adult brain. Scale bar, 50 μm. A-P, anterior to posterior; L-R, left-right; OB, olfactory bulb.

### Aversive olfactory cues activate *lratd2a* neurons in the right dHb

Several studies using transgenic expression of the genetically encoded calcium indicator GCaMP have demonstrated that dHb neurons respond to olfactory cues in larval zebrafish (Jetti *et al*., 2014; Krishnan *et al*., 2014). To determine whether olfactants activate the *lratd2a* positive neuronal cluster in the right dHb, we analyzed calcium signaling in *Tg(lratd2a:QF2)*^*c644*^, *Tg(QUAS:GCaMP6f)*^*c587*^ larval and juvenile zebrafish. Following exposure of 7, 14 and 21 dpf individuals to cadaverine (Figures 2A-B, Supplementary figures 2 and 3), a known aversive olfactory cue that is released from decaying fish (Hussain et al., 2013), we measured a two-fold increase in activity compared to that after application of water alone. The *lratd2a* dHb neurons also responded to chondroitin sulfate, a component of alarm substance (also known as Schreckstoff), which is released from the skin of injured fish (Mathuru *et al*., 2012). A two-fold increase in calcium signals was detected after the first and second delivery of chondroitin sulfate to 22 dpf larvae and a four-fold increase after the second application to 7 and 14 dpf larvae (Figures 2A-B, Supplementary figures 2 and 3).

**Figure 2.**
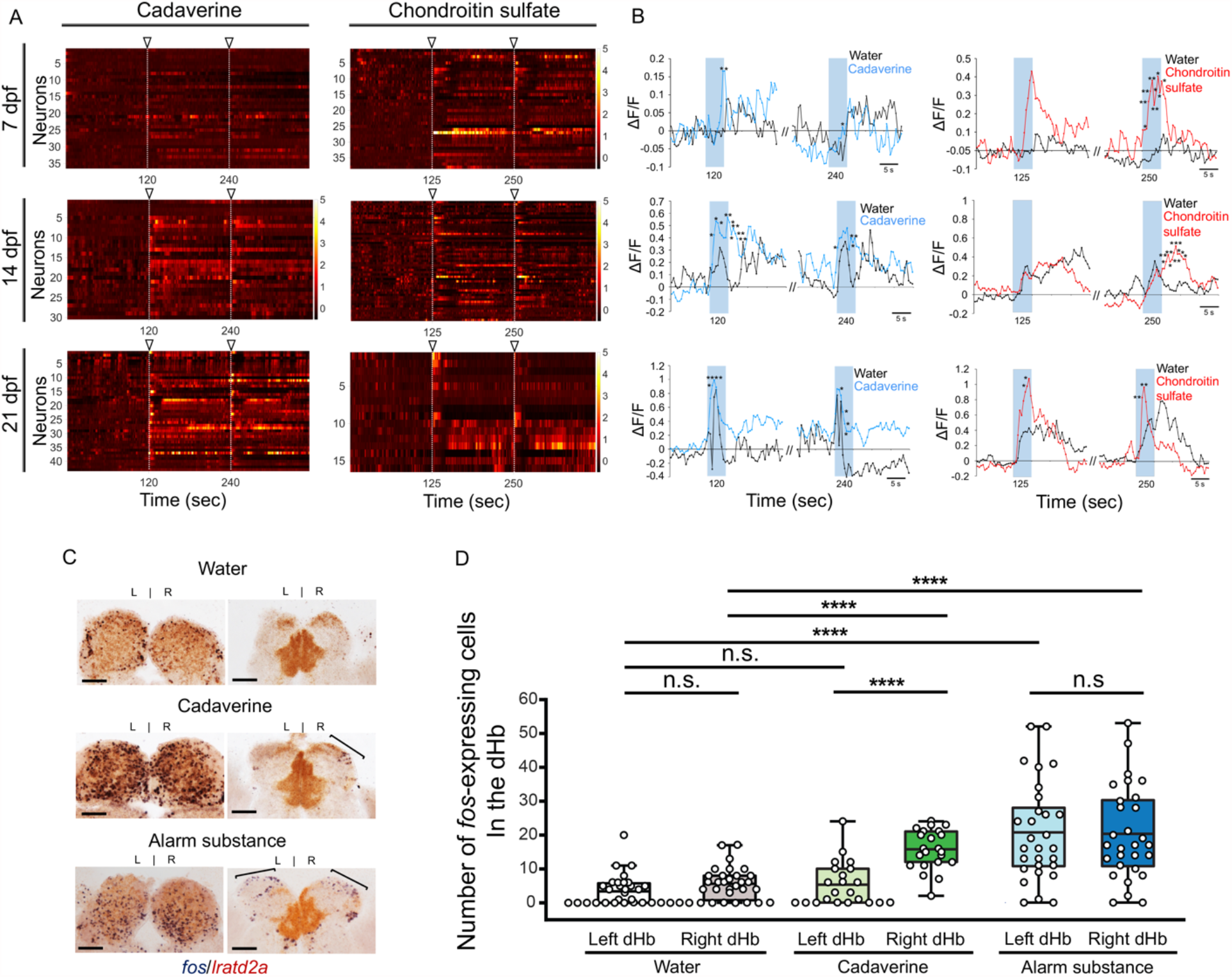
Increased activity of *lratd2a*-expressing dHb neurons upon exposure to aversive olfactory cues. (A) Change in GCaMP fluorescence intensity (ΔF/F) of *lratd2a* neurons in the right dHb in response to cadaverine (*n* = 36 neurons in 5 larvae at 7 dpf, 30 neurons in 5 larvae at 14 dpf and 43 neurons in 5 larvae at 21 dpf) or chondroitin sulfate (*n* = 38 neurons in 4 larvae at 7 dpf, 53 neurons in 5 larvae at 14 dpf and 16 neurons in 2 larvae at 21 dpf). Open arrowheads indicate the time of odorant delivery. (B) Average change in fluorescence (ΔF/F) in seconds (sec) for all *lratd2a* positive neurons in the larval right dHb. Blue bars indicate 5 sec periods of odor delivery. (C) *fos* (blue) and *lratd2a* (brown) transcripts in adult olfactory bulbs (left panels) and habenulae (right panels) detected by RNA *in situ* hybridization 30 min after addition of water, cadaverine or alarm substance to the test tank. Brackets indicate *fos-*expressing cells. Scale bar, 100 μm. (D) Quantification of *fos*-expressing cells in the adult dHb after addition of vehicle alone [3.58 ± 0.92 cells in the left and 5.61 ± 1.07 in the right dHb, *n* = 16 fish (Mann-Whitney *U* = 349, *P* = 0.065)], cadaverine [5.32 ± 1.36 cells in the left and 15.73 ± 1.25 in the right dHb, *n* = 11 fish (Mann-Whitney *U* = 57, P<0.00001)], or alarm substance [20.72 ± 2.70 cells in the left and 20.31 ± 2.53 in the right dHb, *n* = 17 fish (Mann-Whitney *U* = 414.5, *P* = 0.928)]. For the right dHb, significantly more cells were *fos* positive after addition of either cadaverine (*P* = 5.5217E-08) or alarm substance (*P* = 3.78292E-06). For the left dHb, a significant difference was only observed after addition of alarm substance (P=7.44458 E-07). For B and D, Student’s t-test. \**P* < 0.05; \*\**P* < 0.01; \*\*\**P* < 0.001; \*\*\*\**P* < 0.0001; *n*.*s*., not significant (P > 0.05).

**Figure 3.**
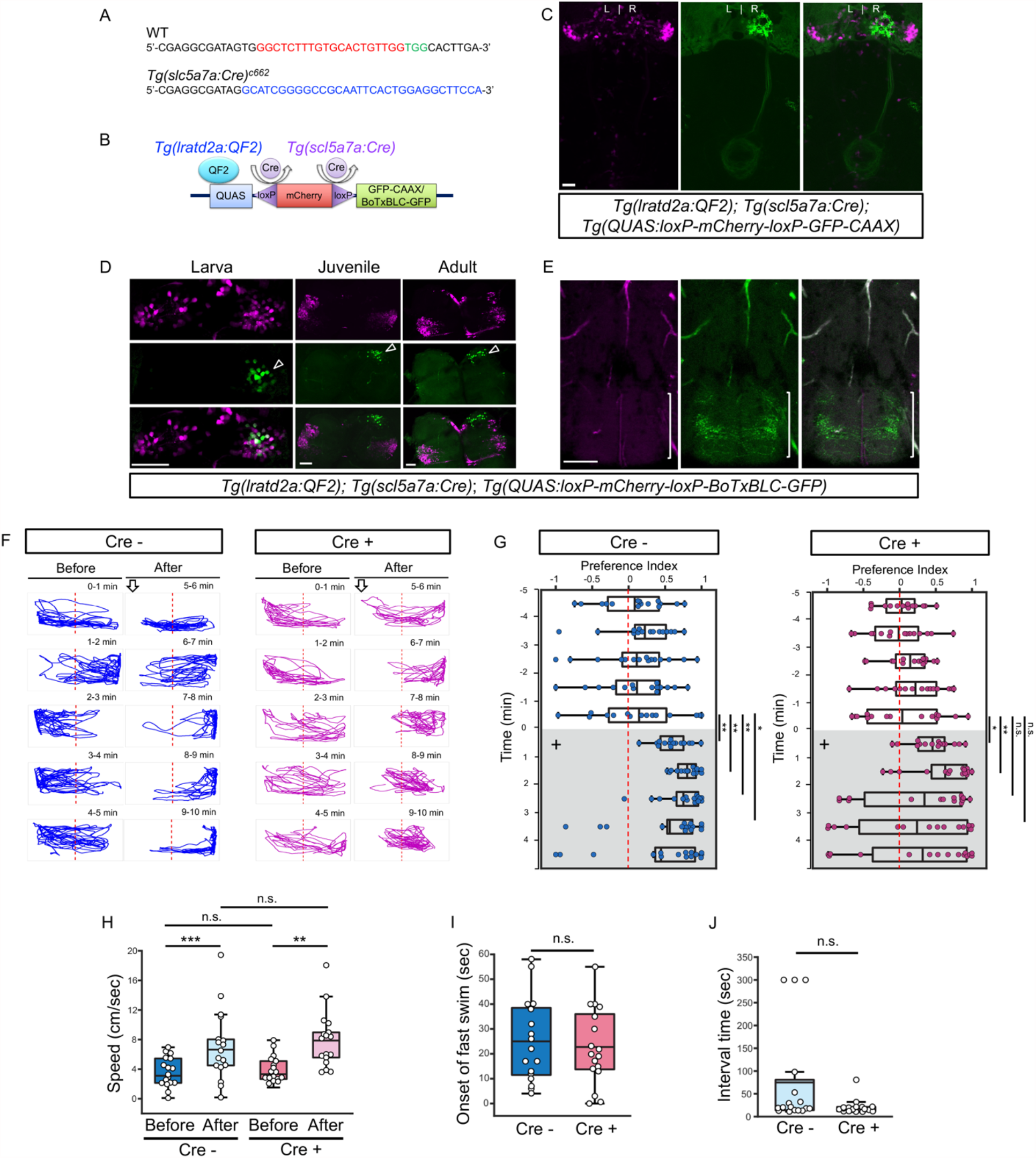
Synaptic inhibition of *lratd2a* right dHb neurons attenuates response to cadaverine. (A) Sequences upstream of the *slc5a7a* transcriptional start site before (WT) and after integration of Cre (blue indicates donor DNA) at sgRNA target site (sgRNA binding site red and PAM sequences green). (B) Schematic diagram of intersectional strategy using Cre/lox mediated recombination and the QF2/QUAS binary system. QF2 is driven by *lratd2a* regulatory sequences and the *slc5a7a* promoter drives Cre leading to reporter/effector expression in *lratd2a* neurons in the right dHb. (C) Dorsal view of GFP labeling in only the right dHb after Cre-mediated recombination in a 5 dpf *Tg(lratd2a:QF2), Tg(slc5a7a:Cre), Tg(QUAS:loxP-mCherry-loxP-GFP-CAAX)* larva. Scale bar, 25 μm. (D) *BoTxBLC-GFP* labeled cells (open arrowhead) in the right dHb in *Tg(lratd2a:QF2), Tg(slc5a7a:Cre), Tg(QUAS:loxP-mCherry-loxP-BoTxBLC-GFP)* 5 dpf, 37 dpf and 4 mpf zebrafish. Upper images show mCherry labeled *lratd2a* Hb neurons, middle images show the subset of right dHb neurons that expressed Cre and switched to GFP, and the bottom row are merged images. Scale bar, 50 μm. (E) Transverse section of *BoTxBLC-GFP* labeled axonal endings of dHb neurons that express Cre and *lratd2a* in a subregion of the vIPN (bracket) in 37 dpf *Tg(lratd2a:QF2), Tg(slc5a7a:Cre), Tg(QUAS:loxP-mCherry-loxP-BoTxBLC-GFP)* juveniles. Scale bar, 50 μm. (F, G) Preferred tank location prior to and after cadaverine addition of adults genotyped for absence (Cre-, blue) or presence (Cre+, red) of *Tg(slc5a7a:Cre)*. (F) Representative 1 min traces for single Cre- and Cre+ adults recorded over 10 mins prior to (mins 0-5) and after (mins 6-10) addition of cadaverine to end of test tank (arrows). (G) Preference index for all adults tested 5 min prior to (white) and 5 min after (grey) addition of cadaverine (on + side). In Cre-fish, significant differences in avoidance behavior were detected after addition of cadaverine [6 min (P=0.0075), 7 min (P=0.0011), 8 min (P=0.0019), 9 min (P=0.032) compared to last min before addition, Student’s t-test, *n* = 15 fish]. Cre+ fish, showed no significant differences in their preferred location beyond two mins after cadaverine addition [6 min (P=0.012), 7 min (P=0.0029) compared to last min before addition, Student’s t-test, n = 15 fish]. Dashed red lines in F and G denote midpoint of test tank. (H) Swimming speed (cm/sec) during 1 min before and after addition of alarm substance for Cre-[3.61 ± 0.48 and 7.13 ± 1.15 , Student’s t-test (P=0.00097)], and Cre+ [4.02 ± 0.42 and 7.93 ± 0.92, Student’s t-test (P=0.001)] adults. (I) Onset of fast swimming after application of alarm substance was observed at 25 ± 4.05 sec in Cre- and at 22.7 ± 3.72 sec in Cre+ fish. (J) Time interval between increased swimming speed and freezing behavior for Cre-(75 ± 26.53 sec) and for Cre+ (20.88 ± 3.93 sec). For H-J, numbers represent the mean ± SEM for *n* = 17 fish. *P < 0.05; **P < 0.01; ***P < 0.001; ****P < 0.0001; *n*.*s*., not significant (P > 0.05).

To examine whether the response to aversive odorants persists in the olfactory-dHb pathway of adult zebrafish, we used expression of the *fos* gene as an indicator of neuronal activation (deCarvalho *et al*., 2013; Hong *et al*., 2013). Consistent with previous findings (Dieris et al., 2017), cadaverine broadly activated OB mitral neurons in the dorsal glomerulus (dG), dorso-lateral glomerulus (dlG), medio-anterior glomerulus (maG), medio-dorsal glomerulus (mdG), lateral glomerulus (lG). In addition, we observed a 3-fold increase in the number of *fos-*expressing neurons in the right dHb following exposure of adult zebrafish to cadaverine relative to delivery of water alone (15.73 ± 1.25 vs. 5.61 ± 1.07 cells, Figures 2C-D). The position of the *fos*-expressing cells in the right dHb corresponds to that of the *lratd2a*-expressing neurons (Figure 2C). Thus, the *lratd2a* subpopulation in the right dHb responds to cadaverine in both larvae and adults.

Exposure to alarm substance prepared from adult zebrafish increased the number of *fos*-expressing mitral cells not only in the lateral glomerulus (lG) and dlG of the OB as would be expected (Mathuru *et al*., 2012; Yoshihara, 2014), but also in the dorso-lateral region of the dHb (Figures 2C-D). In contrast to cadaverine, alarm substance activated neurons in both the left and right dHb (20.72 ± 2.70 cells on the left and 20.31 ± 2.53 on the right).

### Synaptic inhibition of right dHb *lratd2a* neurons reduces aversive response to cadaverine

To confirm that *lratd2a* expressing neurons play a role in processing of aversive olfactory cues, we inhibited synaptic transmission in these cells by mating *Tg(lratd2a:QF2)* fish to a newly generated transgenic line, *Tg(QUAS:BoTxBLC-GFP)*^*c605*^, in which Botulinum toxin light chain C (*BoTxBLC*) (Lal *et al*., 2018; Sternberg *et al*., 2016; Zhang *et al*., 2017) is placed under QUAS control. We tested how adults from the *BoTxBLC-GFP* line reacted to cadaverine by introducing the odorant to one end of a test tank, and measuring the time individuals spent within or outside of this region of the tank. Adults bearing *Tg(lratd2a:QF2)* and *Tg(QUAS:BoTxBLC-GFP)* did not actively avoid the end of the tank where cadaverine was introduced (Supplementary figure 4) as did their sibling controls.

The *Tg(lratd2a:QF2)* driver line is expected to inhibit *lratd2a*-expressing neurons in both the vHb as well as in the right dHb. We therefore devised an intersectional strategy that combines Cre/lox mediated recombination (Förster et al., 2017; Satou et al., 2013; Tabor et al., 2019) and the QF2/QUAS system (Ghosh and Halpern, 2016; Subedi *et al*., 2014) to block the activity of neurons selectively in the right dHb. We produced transgenic fish expressing Cre recombinase under the control of the endogenous *solute carrier family 5 member 7a* (*slc5a7a*) gene using CRISPR/Cas9 targeted integration (Kimura *et al*., 2014) (Figures 3A-B). *slc5a7a* encodes a choline transporter involved in acetylcholine biosynthesis and, in zebrafish larvae, is strongly expressed in the right dHb and not in the vHb (Hong *et al*., 2013). Accordingly, in larvae bearing the three transgenes *Tg(lratd2a:QF2)*^*c601*^, *Tg(slc5a7a:Cre)*^*c662*^ and *Tg(QUAS:loxP-mCherry-loxP-GFP-CAAX*)^*c679*^ Cre-mediated recombination resulted in a switch in reporter labeling from red to green in right dHb neurons (Figure 3C). We followed a similar approach to inhibit synaptic transmission from *lratd2a* right dHb neurons using Botulinum neurotoxin (Lal *et al*., 2018; Sternberg *et al*., 2016; Zhang *et al*., 2017). A *BoTxBLC-GFP* fusion protein was placed downstream of a floxed *mCherry* reporter to make *Tg(QUAS:loxP-mCherry-loxP-BoTxBLC-GFP)*^*c674*^. To validate the effectiveness of this transgenic line, a neuron specific promoter from a Xenopus neural-specific beta tubulin (*Xla*.*Tubb2*) gene (Peri and Nusslein-Volhard, 2008) was used to drive QF2 expression. Larvae bearing *Tg(Xla*.*Tubb2:QF2;he1*.*1:mCherry)*^*c663*^; *Tg(slc5a7a:Cre)*^*c662*^ and *Tg(QUAS:loxP-mCherry-loxP-BoTxBLC-GFP)*^*c674*^ showed a significantly reduced response to a touch stimulus, indicating that the neurotoxin was produced in the presence of Cre recombinase (Supplementary figure 5, Supplementary movie 1).

*BoTxBLC-GFP* was selectively expressed in *lratd2a*/*slc5a7a* neurons of the right dHb (Figure 3D and Supplementary figure 6) in larvae bearing the three transgenes *Tg(lratd2a:QF2)*^*c601*^, *Tg(slc5a7a:Cre)*^*c662*^ and *Tg(QUAS:loxP-mCherry-loxP-BoTxBLC-GFP)*^*c674*^. Axons labeled by *BoTxBLC-GFP* terminated at the vIPN in the same location as those observed in *Tg(lratd2a:QF2)* (Figure 3E), suggesting that botulinum neurotoxin inhibits synaptic transmission within this restricted region of the vIPN.

To determine whether the *lratd2a* neurons in the right dHb contributed to the aversive response to cadaverine, we monitored behavior following its addition . During the first two minutes following exposure, adults both expressing or not expressing *BoTxBLC-GFP* avoided cadaverine. However, the aversive response was sustained for 4 min in control fish, but not in those expressing *BoTxBLC-GFP* in the *lratd2a* neurons of the right dHb (Figures 3F-G). These findings indicate that this subset of right dHb neurons are required for a prolonged aversive response to cadaverine.

Disruption of synaptic transmission in *lratd2a*-expressing Hb neurons alone did not alter the response of zebrafish to alarm substance, which typically triggers erratic, rapid swimming and bottom dwelling, followed by freezing behavior (Diaz-Verdugo et al., 2019; Jesuthasan and Mathuru, 2008). Similar to controls, both juveniles and adults expressing *BoTxBLC-GFP* under the control of *Tg(lratd2a:QF2)*^c601^ showed rapid swimming/darting behavior within 22-25 second after delivery of alarm substance, first doubling their speed of swimming (Figures 3H-J and Supplementary figure 7), and then freezing for the duration of the 5 min recording period Blocking the activity of *lratd2a* neurons in the right dHb and in the bilateral vHb is therefore insufficient to diminish the robust behavioral changes elicited by alarm substance (Figures 3H-J and Supplementary figures 4C-D).

### Zebrafish mutants with habenular defects show altered responses to aversive cues

We examined the response to aversive odorants by *tcf7l2*^*zf55*^ mutants that develop with symmetric left-isomerized dHb and lack the vHb, but are viable to adulthood (Husken et al., 2014). In agreement with the transformation of dHb identity, projections from OB mitral cells do not terminate in the right dHb of homozygous mutants nor are *lratd2a*-expressing neurons or their efferents to the vIPN detected (Figures 4A-D).

**Figure 4.**
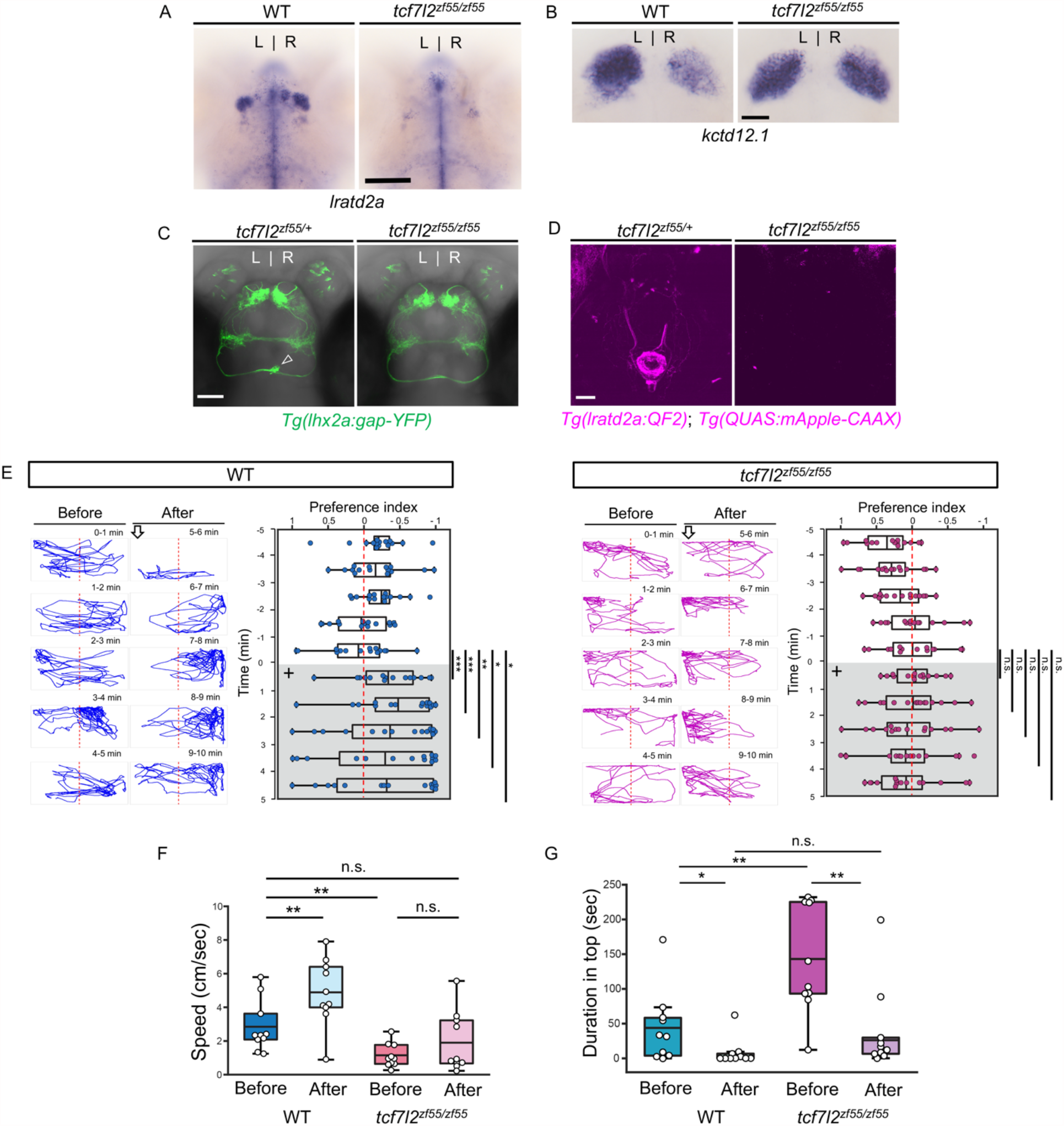
Attenuated response to aversive odorants in left-isomerized dHb mutants. (A-B) (A) Absence of *lratd2a*-expressing right dHb neurons and (B) right-isomerized expression of *kctd12*.*1* in *tcf7l2* mutant larvae at 5 dpf. (C) Dorsal views of olfactory mitral neuronal projections of *Tg(lhx2a:gap-YFP)* larvae at 6 dpf. Open arrowhead indicates axon terminals of mitral cells in the WT right dHb that are absent in the mutant. (D) Dorsal views of dHb neuronal projections to the ventral IPN in *Tg(lratd2a:QF2), Tg(QUAS:mApple-CAAX)* larvae at 6 dpf. (E) Representative traces (1 min) and preference index for *tcf7l2* mutant and WT sibling adults after addition of cadaverine (on + side). Only the WT siblings showed a significant difference in avoidance behavior for the 5 mins afterwards compared to the min before its addition [6 min (P=0.00035), 7 min (P=0.000175), 8 min (P=0.0041), 9 min (P=0.0177), 10 min (P=0.0203), *n* = 15 adults. Student’s t-test]. (F) Swimming speed (cm/sec) for 30 sec before and after addition of alarm substance. For *tcf7l2* homozygotes, 1.13 ± 0.22 cm/s and 1.89 ± 0.56 cm/s [Student’s t test (P = 0.1011)] and for their WT siblings 2.84 ± 0.48 cm/s and 4.88 ± 0.63 cm/s [Student’s t test (P=0.0026)], *n* = 10 fish for each group (G) Duration in the upper half of the test tank prior to and after addition of alarm substance for *tcf7l2* adults was 143.58 ± 24.80 sec and 38.77 ± 19.56 sec (Mann-Whitney U=12; P=0.0047) and for their WT siblings was 43.68 ± 16.35 sec and 8.19 ± 6.16 sec (Mann-Whitney U=20.5, P=0.0285), *n* = 10 fish for each group. For F-G, numbers represent the mean ± SEM. \**P* < 0.05; \*\**P* < 0.01; \*\*\**P* < 0.001; *n*.*s*., not significant (P > 0.05).

Following application of cadaverine, *tcf7l2*^*zf55*^ homozygous adults did not exhibit the characteristic avoidance behavior of their wild type siblings (Figure 4E). Exposure to alarm substance also did not elicit a significant increase in their swimming speed from baseline (1.13 ± 0.22 cm/sec before and 1.89 ± 0.56 cm/sec after) relative to WT siblings (2.84 ± 0.48 cm/sec before and 4.88 ± 0.63 cm/sec after, Figure 4F). However, homozygous *tcf7l2*^*zf5*^ mutants tended to spend more time swimming in the top half of a novel test tank than wild-type adults, a behavior that was suppressed in the presence of alarm substance (Fig. 4G).

To further assess the role of *lratd2a*-expressing neurons in aversive olfactory processing, we looked at homozygous mutants of the *brain-specific homeobox* (*bsx*) gene, which develop right-isomerized dHb [(Schredelseker and Driever, 2018) and Figure 5A] and are viable to adulthood (Schredelseker and Driever, 2018). As might be expected when both dHb have right identity, equivalent populations of *lratd2a-*expressing neurons were found on both sides of the brain (Figure 5C). Instead of innervating only the right dHb as in controls, the axons of *lhx2a*:*gap*-*YFP* labeled olfactory mitral cells terminated in the left and right dHb [(Dreosti *et al*., 2014) and Figure 5B], where the clusters of *lratd2a* neurons are situated (data not shown). Projections from the *lratd2a* dHb neurons coursed bilaterally through the left and right fasciculus retroflexus (FR) and innervated the same limited region of the ventral IPN (Figure 5C).

**Figure 5.**
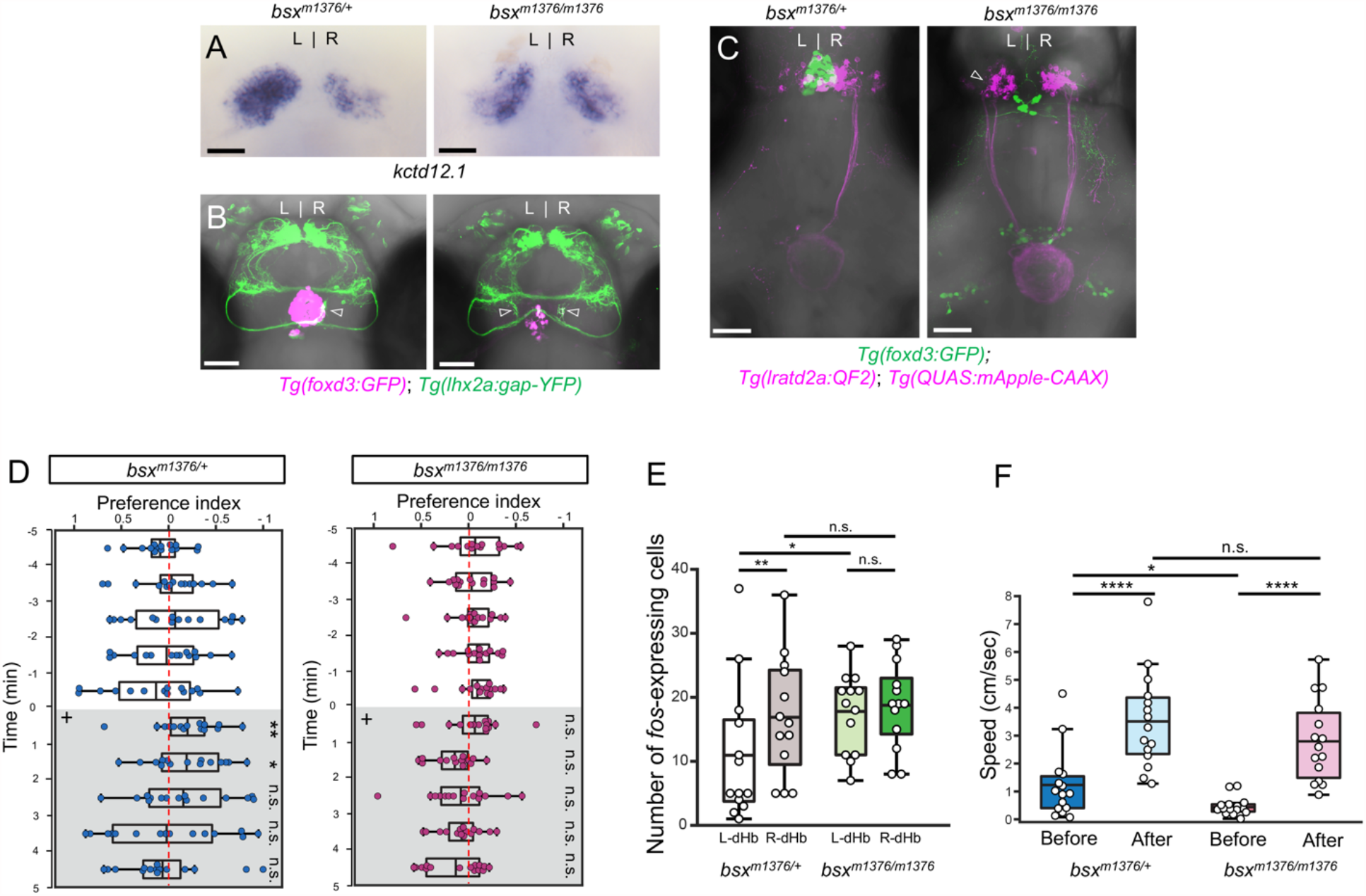
Enhanced reactivity to alarm substance in mutants with right-isomerized dHb. (A) Asymmetric expression of *kctd12*.*1* is right-isomerized in *bsx* homozygotes at 5 dpf. (B) Projections of *Tg(lhx2a:gap-YFP)* labeled olfactory mitral cells terminate bilaterally (open arrowheads) at *lratd2a* neurons in *bsx*^*m1376*^ homozygous mutants at 5 dpf. (C) In the mutants, axons from both left (open arrowhead) and right dHb *lratd2a* neurons project to the same region of the vIPN. Scale bar, 50 μm. (D) Preferred tank location of *bsx*^*m1376*^ adults after addition of cadaverine (on + side). Only the heterozygotes showed a significant difference in preference after application of cadaverine compared to the min before its addition [6 min (Mann-Whitney *U* = 37; *P* = 0.00188), 7 min (Mann-Whitney *U* = 56; *P* = 0.02034). *n* = 15 adults for each group]. (E) Quantification of *fos*-expressing cells in the dHb after application of cadaverine in *bsx*^*m1376/+*^ [11 ± 2.97 cells on the left and 16.84 ± 2.62 cells on the right. Student’s t-test (P=0.006)] and *bsx*^*m1376/m1376*^ adults [17.61 ± 1.72 cells on the left and 18.77 ± 1.88 cells on the right. Student’s t-test (P=0.645)], for *n* = 13 sections from 7 adults for each group. (F) Swimming speed (cm/sec) for 30 sec before and after addition of alarm substance. In heterozygous adults, swimming speed was 1.22 ± 0.31 cm/s before and 3.52 ± 0.44 cm/s after [Student’s t test (P=9.22089E-05)] and in homozygotes, 0.46 ± 0.08 cm/s before and 2.80 ± 0.37 cm/s after [Student’s t test (P=1.38481E-05)], *n* = 15 adults for each group. For E-F, numbers represent the mean ± SEM. \**P* < 0.05; \*\**P* < 0.01; \*\*\*\**P* < 0.0001; *n*.*s*., not significant (P > 0.05).

To measure the reaction to cadaverine in *bsx*^*m1376*^ homozygous adults with bilaterally symmetric *lratd2a* neurons, we counted the number of cells expressing *fos* in the dHb and found an increase in the left nucleus compared to heterozygous siblings (Figure 5E, Supplementary figure 8). Despite the symmetric activation of dHb neurons, *bsx*^*m1376/m1376*^ mutants did not exhibit increased avoidance to cadaverine compared to controls (Figure 5D). Overall, homozygous mutants were slower swimmers than heterozygotes; however, after exposure to alarm substance, their swimming speed relative to baseline was two-fold faster than that of their heterozygous siblings (Figure 5F), indicative of an enhanced response to this aversive cue.

## Discussion

From worms to humans, stimuli including odors are differently perceived by left and right sensory organs to elicit distinct responses (Gunturkun and Ocklenburg, 2017; Güntürkün et al., 2020). Honeybees, for example, show an enhanced performance in olfactory learning when their right antenna is trained to odors (Guo et al., 2016; Letzkus et al., 2006; Rogers and Vallortigara, 2008). In mice, over one third of mitral/tufted cells were found to be interconnected between the ipsilateral and contralateral olfactory bulbs for sharing of odor information received separately from each nostril, and for coordinated perception (Grobman et al., 2018). The zebrafish provides a notable example of a lateralized olfactory pathway, with the discovery of a subset of bilateral mitral cells that project to the dorsal habenulae but terminate only at the right nucleus (Miyasaka *et al*., 2014; Miyasaka *et al*., 2009). This finding prompted us to ask what is different about the post-synaptic dHb neurons that receive this olfactory input and what function does this asymmetric pathway serve.

### Aversive olfactory cues activate identified neurons in the right dHb

We previously showed that the olfactory mitral cells that express *lhx2a* and are located in medio-dorsal and ventro-medial bilateral glomerular clusters (Miyasaka *et al*., 2014; Miyasaka *et al*., 2009) project their axons to a subregion of the right dHb where the *lratd2a* gene is transcribed (deCarvalho *et al*., 2013). From transgenic labeling with membrane-tagged fluorescent proteins, we now confirm that the *lhx2a* olfactory neurons precisely terminate at a cluster of *lratd2a*/*slc5a7a* expressing cholinergic neurons present in the right but not the left dHb.

From calcium imaging, we validated that the right dHb appears more responsive than the left when larval zebrafish are exposed to aversive odors such as cadaverine or chondroitin sulfate (Jetti *et al*., 2014; Krishnan *et al*., 2014), a component of alarm substance (Mathuru *et al*., 2012), and further determined that the *lratd2a*-expressing neurons of the right dHb specifically respond to these aversive olfactory cues. As has also been observed by others (Jesuthasan et al., 2020), application of vehicle alone, even when introduced slowly into a testing chamber, is sufficient to elicit a change in GCaMP fluorescence. Determining the habenular response to odorants relative to vehicle alone is thus an essential measure, but one that has not been reported in all studies (Chen *et al*., 2019; Jetti *et al*., 2014; Krishnan *et al*., 2014).

In adults, we used *fos* expression as a measure of neuronal activation and showed that transcripts colocalized to *lratd2a*-expressing cells. Interestingly, cadaverine predominantly activated neurons in the right dHb in larvae and adults, whereas neurons responsive to alarm substance were detected in both the left and right dHb nuclei of adult zebrafish. Different types of olfactory cues activate distinct glomeruli in the OB (Friedrich and Korsching, 1997; Yoshihara, 2014), and consistent with the prior studies, we observed that, in adults, cadaverine significantly increased *fos* expression in the mdG and dG regions, the location of *lhx2a* neurons that project to the right dHb. By contrast, alarm substance predominantly activated neurons in the lG and dlG regions of the OB that innervate the telencephalon and posterior tuberculum (Miyasaka *et al*., 2014; Miyasaka *et al*., 2009), suggesting that both dHb receive input via this route rather than through direct olfactory connections. Indeed, we found that more neurons reacted to alarm substance than cadaverine throughout the brain, including in the Dp, Vv and thalamic areas (data not shown).

In previous experiments (deCarvalho *et al*., 2013), we did not detect activated neurons in the right dHb of adult zebrafish following exposure to cadaverine or alarm substance. Several factors could account for the difference from the earlier study: we now have the transgenic tools to examine *lratd2a* neurons directly, we used higher concentrations of cadaverine and alarm substance and, in contrast to delivering odorants to groups of zebrafish, we tested the neuronal response in individual adults.

It has been suggested that lateralized olfactory and visual functions of the dHb are more prominent early in development and less so at later stages (Fore et al., 2020). However, the presence of *lratd2a*-expressing neurons in the right dHb and their preferential response to cadaverine from larval to juvenile and adult stages supports the persistence of lateralized activity and illustrates the value of examining defined neuronal populations.

### Right dHb neurons mediate aversive behavioral responses

Despite both being aversive cues (Hussain *et al*., 2013; Mathuru *et al*., 2012), cadaverine and alarm substance elicit different behavioral responses by adult zebrafish. Control fish show active avoidance to cadaverine for the first to 2 to 4 minutes of a 5 minute testing period, whereas alarm substance triggers immediate erratic behavior such as rapid swimming and darting that is typically followed by prolonged freezing (Hussain *et al*., 2013; Mathuru *et al*., 2012).

We found that perturbation of the *lratd2a*-expressing right dHb neurons either selectively by *BoTxBLC*-mediated synaptic inactivation, or in *tcf7l2*^*zf55*^ homozygous mutants that completely lack them, reduced avoidance to cadaverine, either in the length or degree of the response.

However, juveniles or adults with *BoTxBLC* inactivated neurons displayed a similar response to alarm substance as controls. In contrast, *tcf7l2*^*zf55*^ mutants, showed no difference in their swimming behavior before and after its addition. One explanation is that many regions throughout the brain are likely involved in directing the complex repertoire of behaviors elicited by alarm substance and inactivation of *lratd2a* neurons in the habenular region alone is insufficient to weaken the overall response. Our findings also rule out a role for the ventral habenulae in the response to alarm substance, as the reaction to alarm substance was intact in transgenic adults in which *lratd2a* neurons were inactivated by *BoTxBLC* in the bilateral vHb as well as in the right dHb. Alternatively, the *tcf7l2*^*zf55*^ mutation could disrupt other brain regions that regulate behaviors elicited by alarm substance since the *tfc7l2* gene is expressed in neurons throughout the brain, including the anterior tectum, dorsal thalamus and the hindbrain (Young et al., 2002).

Similar to *tcf7l2*^*zf5*^, the *bsx*^*m1376*^ mutation is pleiotropic not only resulting in right-isomerization of the dHb due to the absence of the parapineal (Schredelseker and Driever, 2018), but also loss of the terminal tuberal hypothalamus, mammillary hypothalamic regions and secondary prosencephalon (Schredelseker et al., 2020). Although homozygous mutants showed a hyperactive response to alarm substance relative to controls, we cannot discount the involvement of other affected brain regions. Albeit technically challenging in adults, a more selective test such as optogenetic activation of only the *lratd2a* dHb neurons in wild-type and mutant zebrafish could help resolve their contribution to the alarm response.

The identification of a subset of neurons in the right dHb that receive olfactory input and terminate their axons at a defined subregion of the ventral IPN lays the groundwork for tracing an entire pathway from olfactory receptors to the neurons directing the appropriate behavioral response. The midline IPN has been morphologically defined into subregions (deCarvalho *et al*., 2014; Lima et al., 2017; Quina et al., 2017), but their connectivity and functional properties have been understudied. Recent work has begun to assign different functions to given subregions, such as the role of the rostral IPN in nicotine aversion (Morton et al., 2018; Quina *et al*., 2017). Neurons in the ventral IPN project to the raphe nucleus (Agetsuma et al., 2010; Lima *et al*., 2017), but the precise identity of raphe neurons that are innervated by the *lratd2a-*expressing dHb neurons remains to be determined. Transcriptional profiling of the IPN should yield useful information on its diverse neuronal populations and likely lead to the identification of the relevant post-synaptic targets in the ventral IPN and their efferent connections. Elaboration of this pathway may also help explain the advantage of lateralization in the processing of aversive information. It has been argued, for instance, that the antennal specialization to aversive odors in bees is correlated with directed turning away from the stimulus and escape (Rogers and Vallortigara, 2019). Beyond olfaction, left-right asymmetry appears to be a more general feature of stress-inducing, aversive responses as demonstrated for the rat ventral hippocampus (Sakaguchi and Sakurai, 2017) and human pre-frontal cortex, where heightened anxiety also activates more neurons on the right than on the left (Avram et al., 2010; Ocklenburg et al., 2016).

## Acknowledgement

We thank Shin-ichi Higashijima for the *Gbait-hsp70:Gal4* donor plasmid, Wenbiao Chen for *pT3TS nCas9n* plasmid, Koichi Kawakami for the *UAS:zBoTXBLC-GFP* construct, Paul Krieg for the *Xla*.*Tubb* promoter construct, Claire Wyart for *GCaMP6f* plasmid and Wolfgang Driever and Tatjana Piotrowski for providing *bsx*^*m1376*^ and *tcf7l2*^*zf5*^ mutant zebrafish, respectively.

## Author contributions

J-H.C. and M.E.H. conceived of and designed the study and wrote the manuscript. J-H.C. performed all of the experiments. E.D. wrote MATLAB script for analyzing behavioral experiments. M.M. constructed sgRNAs for *lratd2a* and *slc5a7a*. J-M.C. generated Tol2 constructs and transgenic lines. All authors reviewed the manuscript.

