## Supplemental Figures for "Specialized neurons in the right habenula mediate response to aversive olfactory cues"

Supplementary figures 1 to 8

Supplementary movie 1

Supplementary table 1

Key resources table

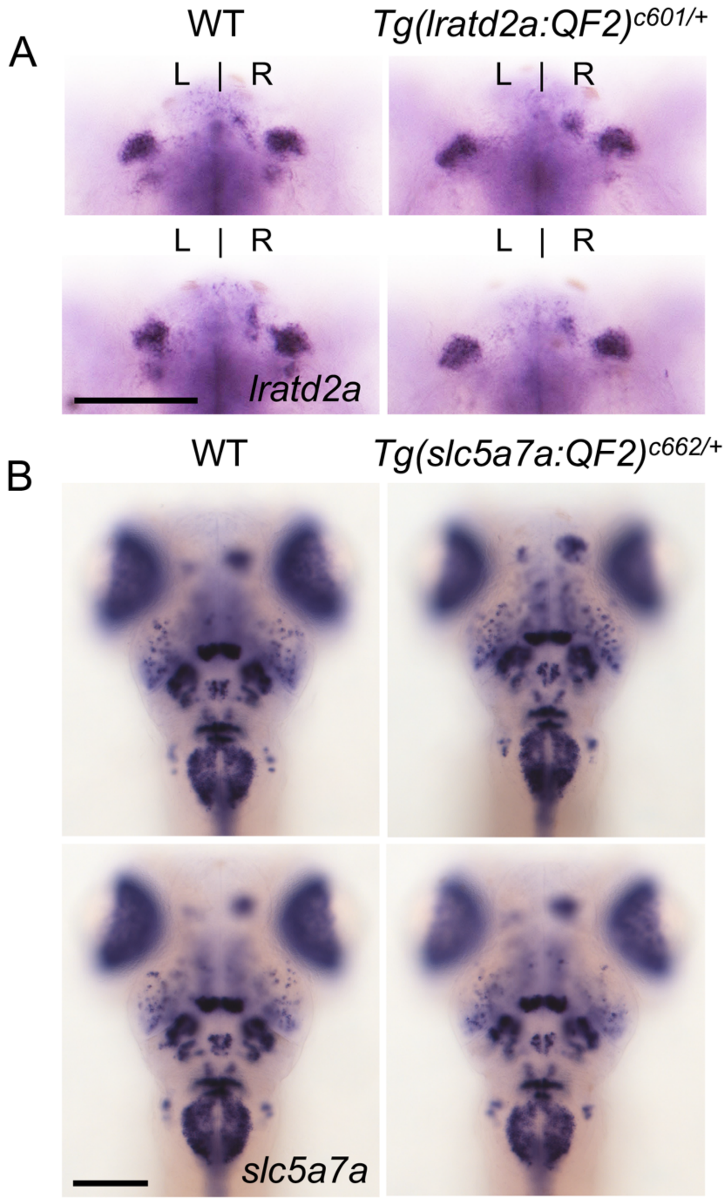


Supplementary figure 1. Targeted genomic integration does not alter endogenous gene expression. Expression patterns of (A) *lratd2a* and (B) *slc5a7a* is similar in 5 dpf wild-type and transgenic larvae. Dorsal views comparing two WT and two transgenic larvae. Scale bar, 100 μm.

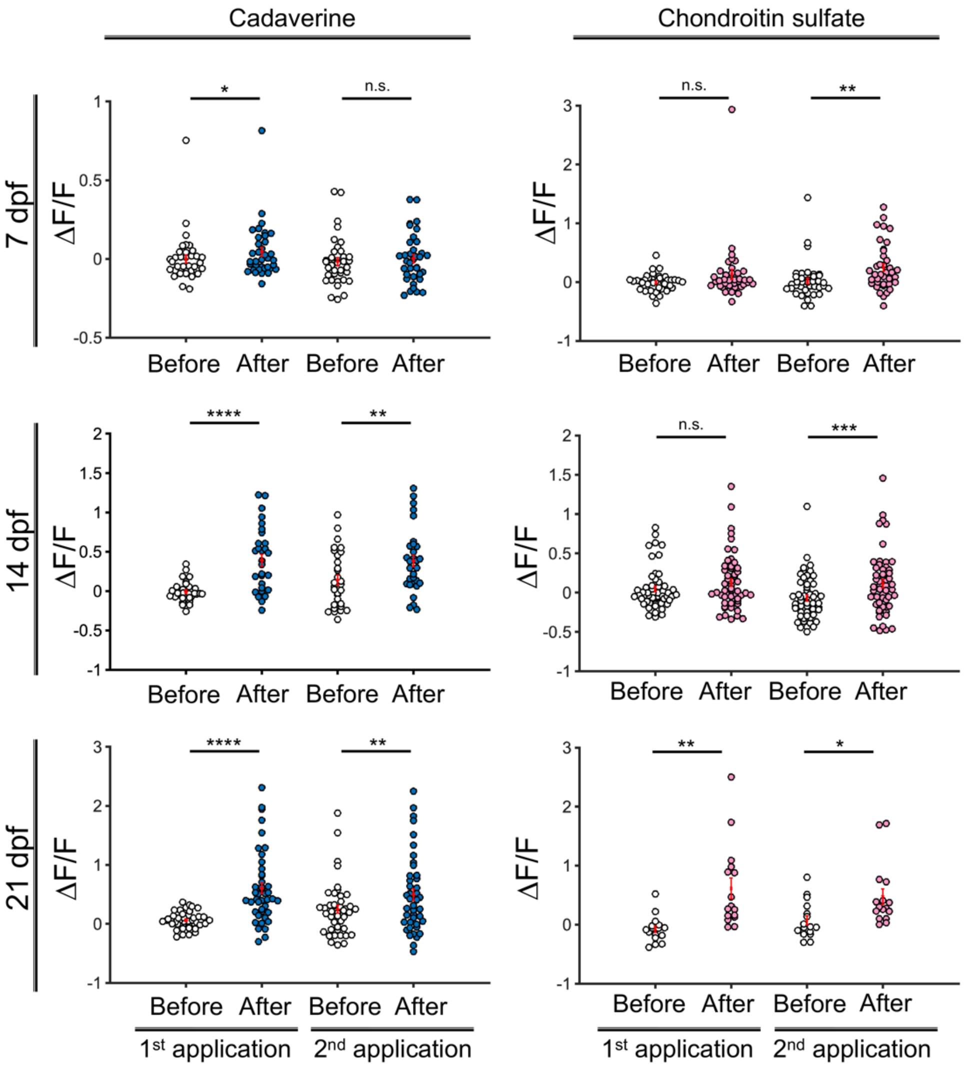


Supplementary figure 2. Activation of identified right habenular neurons following exposure to aversive olfactory cues. Change in GCaMP fluorescence intensity (ΔF/F) in *lratd2a* positive neurons of the right dHb at different developmental stages in response to cadaverine ($n$ = 36 neurons in 5 larvae at 7 dpf, 30 neurons in 5 larvae at 14 dpf and 43 neurons in 5 larvae at 21 dpf) or chondroitin sulfate ($n$ = 38 neurons in 4 larvae at 7 dpf, 53 neurons in 5 larvae at 14 dpf and 16 neurons in 2 larvae at 21 dpf). *$P < 0.05$; **$P < 0.01$; ***$P < 0.001$; ****$P < 0.0001$; *n.s.*, not significant (P > 0.05).

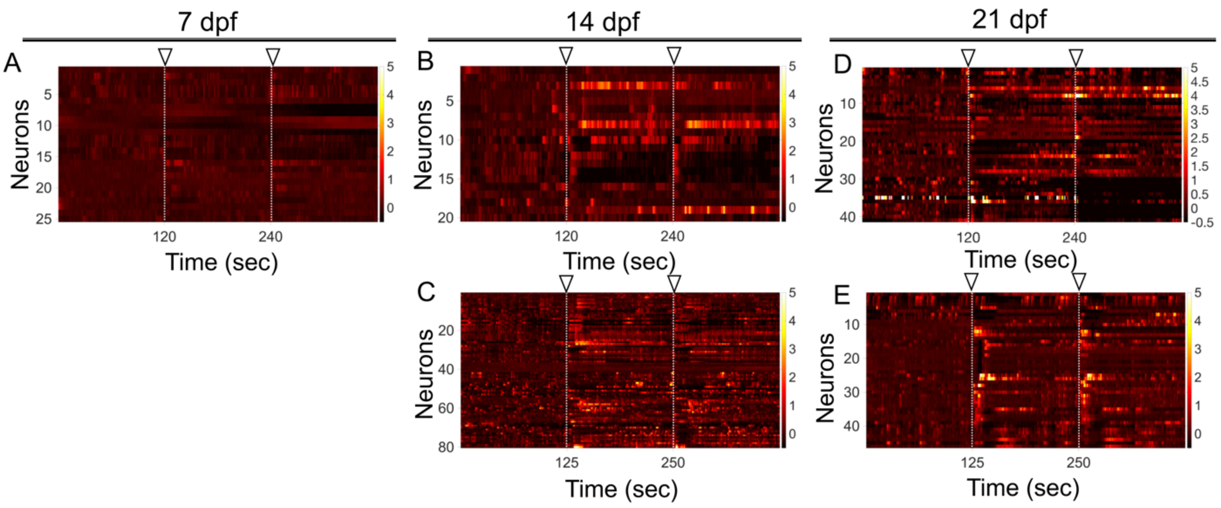


Supplementary figure 3. Response of dHb *lratd2a* neurons to delivery of water. Controls for (A, B, D) cadaverine and (A, C, E) chondroitin sulfate. Open arrowheads indicate time of odorant delivery. At 7 dpf, water control for cadaverine and chondroitin, $n$ = 25 neurons in 5 larvae. At 14 dpf, water control for cadaverine, $n$ = 20 neurons in 4 larvae, and for chondroitin, $n$ = 80 neurons in 7 larvae. At 21 dpf, water control for cadaverine, $n$ = 41 neurons in 5 larvae, and for chondroitin, $n$ = 46 neurons in 7 larvae.

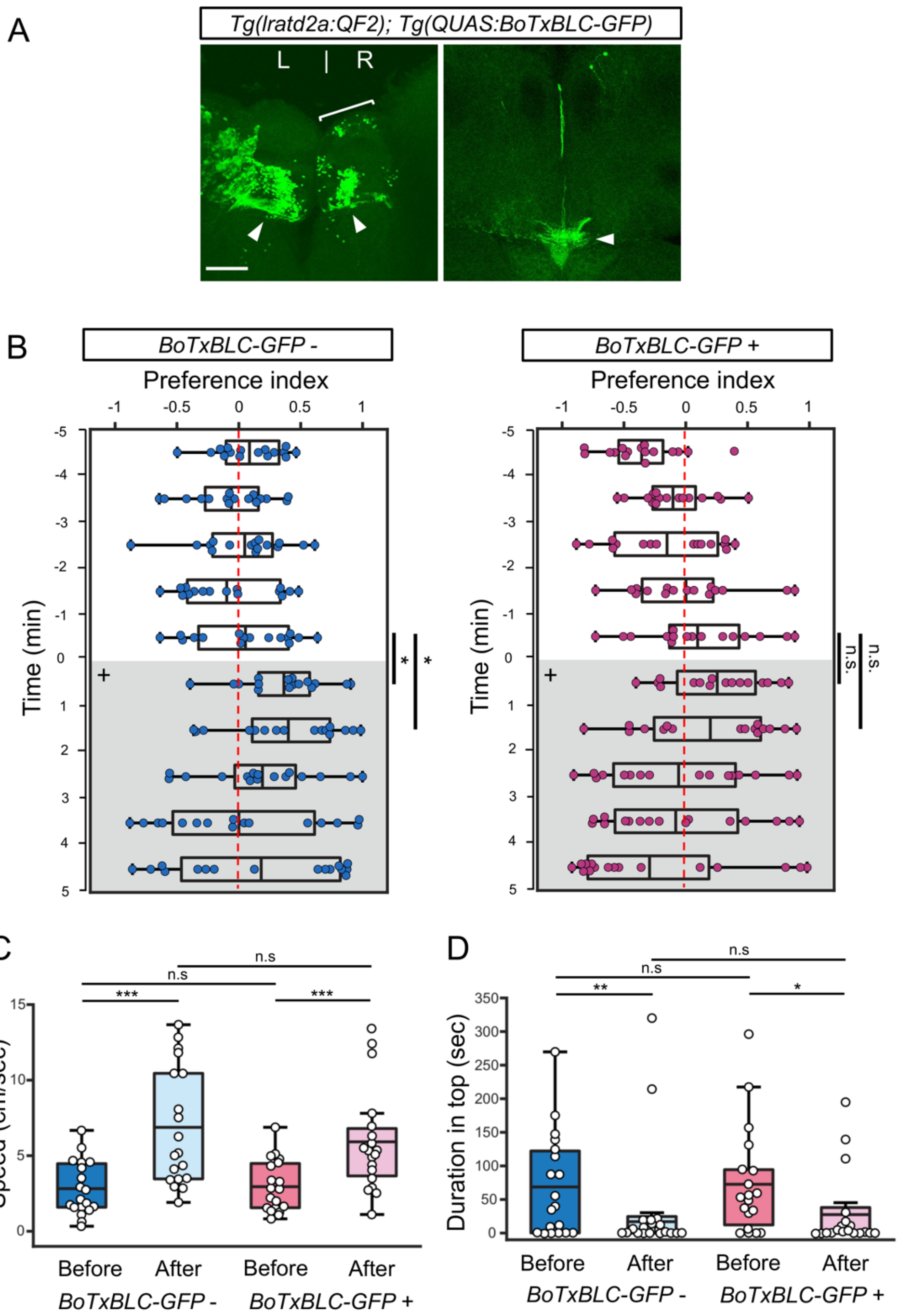


Supplementary figure 4. Synaptic inhibition of *lratd2a*-expressing neurons reduces response to cadaverine. (A) Transverse sections through the Hb and the IPN of a 4 mpf *Tg(lratd2a:QF2)*, *Tg(QUAS:BoTxBLC-GFP)* fish. On the left, the dHb (bracket) and vHb (arrowhead) and on the right the vIPN (arrowhead) are indicated. Scale bar, 100 µm. (B) Preferred tank location of adults genotyped for absence (blue) or presence (red) of *BoTxBLC-GFP* prior to (white) and after (grey) addition of cadaverine (on + side). Adults lacking *BoTxBLC-GFP* showed a significant difference in avoidance behavior compared to the min before its addition [6 min (P=0.015) and 7 min (P=0.019), $n = 16$ fish for each group. Student's t-test]; however, those expressing *BoTxBLC-GFP* in *lratd2a* neurons showed no response to cadaverine. (C) Swimming speed for 1 min before and after addition of alarm substance in fish lacking *BoTxBLC-GFP* was 2.81 ± 0.4 cm/sec and 6.84 ± 0.89 cm/sec, respectively [Student's t-test (P=0.00012)] or expressing *BoTxBLC-GFP*, speed was 2.97 ± 0.39 cm/sec and 5.89 ± 0.77 cm/sec [Student's t-test (P=0.00087)], $n = 19$ fish for each group. (D) Duration in the top half of the test tank before and after addition of alarm substance. Adults lacking the transgene spent 68.24 ± 17.6 sec and 17.06 ± 11.09 sec [Student's t-test (P=0.0049)] and *BoTxBLC-GFP* fish spent 73.07 ± 18.25 sec and 27.71 ± 12.85 sec, [ Student's t-test (P=0.01), $n = 19$ fish for each group. For C-D, number represent the mean ± SEM. *$P < 0.05$; **$P < 0.01$; ***$P < 0.001$; *n.s.*, not significant (P > 0.05).

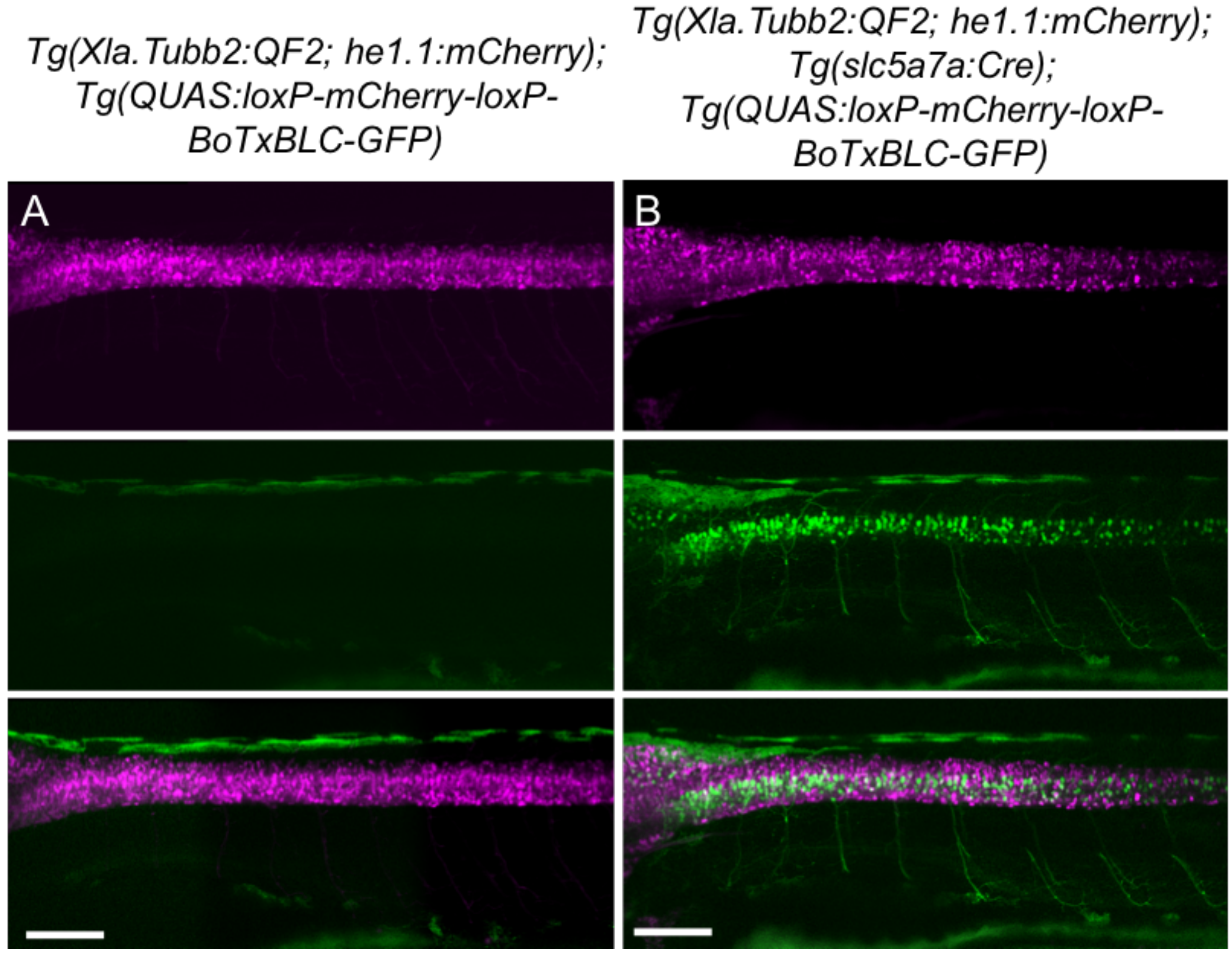


Supplementary figure 5. Validation of intersectional strategy to inhibit cholinergic neurons using a botulinum neurotoxin.

Lateral views of (A) *Tg(Xla.Tubb:QF2)*, *Tg(QUAS:loxP-mCherry-loxP-BoTxBLC-GFP)* and (B) *Tg(Xla.Tubb:QF2)*, *Tg(slc5a7a:Cre)*, *Tg(QUAS:loxP-mCherry-loxP-BoTxBLC-GFP)* larvae at 4 dpf. Scale bar, 100 µm. In the presence of Cre recombinase, cholinergic neurons in the spinal cord switch from mCherry to *BoTxBLC-GFP* expression, which inhibits their response to touch (refer to Supplementary movie 1)

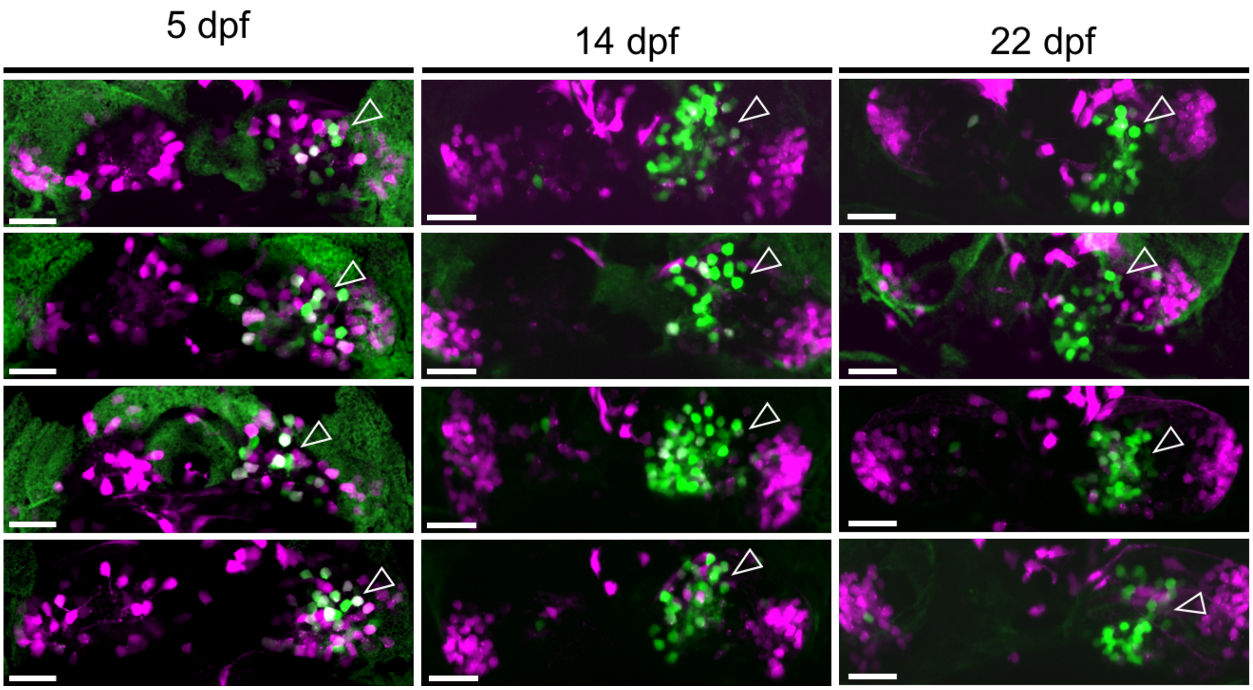


Supplementary figure 6. Variability in *BoTxBLC-GFP* expression.

Dorsal views of *Tg(lratd2a:QF2)*, *Tg(slc5a7a:Cre)*, *Tg(QUAS:loxP-mCherry-loxP-BoTxBLC-GFP)* zebrafish at 5, 14 and 22 dpf showing variability in labeling of *lratd2a* right Hb neurons (open arrowheads). Four individuals are shown for each stage. Scale bar, 25 µm.

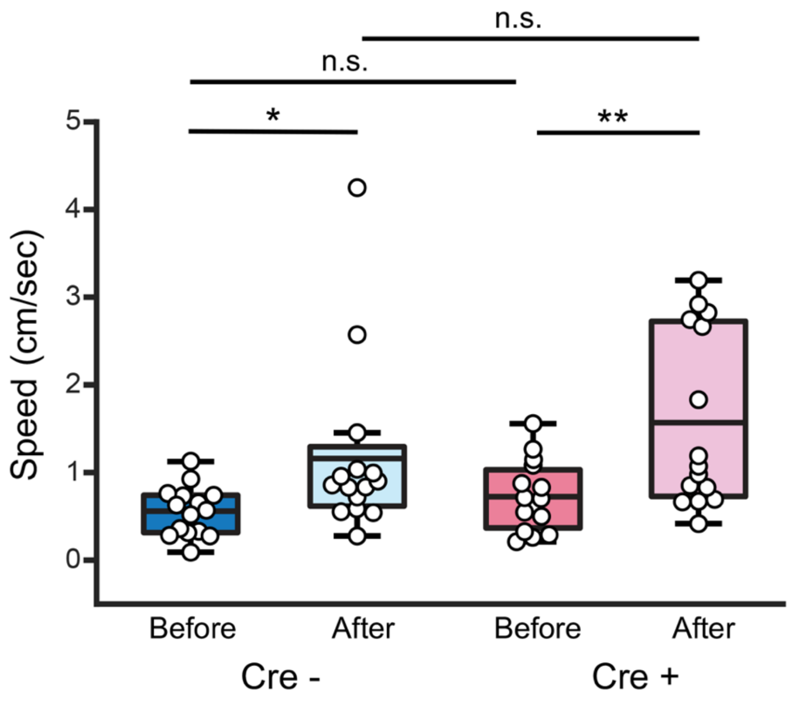


Supplementary figure 7. Aversive response to alarm substance is intact in *BoTxBLC-GFP* juvenile fish. (A) Swimming speed for 1 min before and after addition of alarm substance in *Tg(lratd2a:QF2)*, *Tg(QUAS:loxP-mCherry-loxP-BoTxBLC-GFP* juveniles with or without *Tg(slc5a7a:Cre)*. In the absence of Cre, speed was 0.55 ± 0.07 before and 1.16 ± 0.259 cm/sec after [Student's t-test (P=0.022)] and, in the presence of Cre, speed was 0.73 ± 0.10 before and 1.57 ± 0.260 cm/sec after [Student's t-test (P=0.0016)], $n$ = 15 fish for each group. Numbers represent the mean ± SEM.

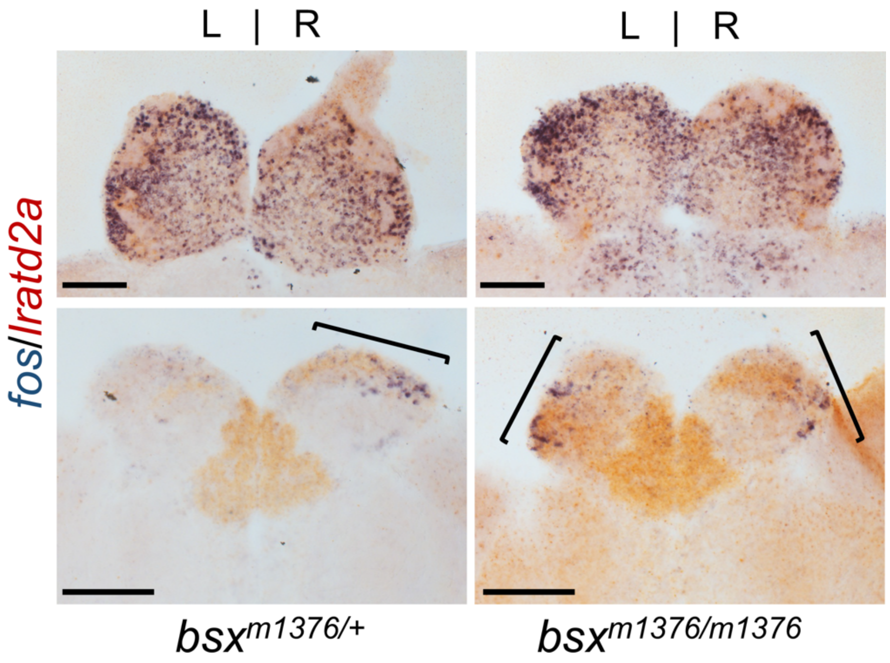


Supplementary figure 8. Bilateral *fos*-expressing neurons in mutants with right-isomerized dHb. *fos* (blue) and *lratd2a* (brown) transcripts in the olfactory bulbs (upper panels) and dHb (bottom panels) of *bsx*$^{m1376}$ heterozygous and homozygous mutants at 10 mpf detected by RNA *in situ* hybridization 30 min after addition of cadaverine to the test tank in. Brackets indicate *fos*-expressing cells. Scale bar, 100 µm

**Supplementary movie 1**

Behavior of *Tg(Xla.Tubb:QF2)*, *Tg(QUAS:loxP-mCherry-loxP-BoTxBLC-GFP)* 4 dpf larvae with or without the *slc5a7a:Cre* transgene in response to touch stimulus.

**Supplementary table 1**

| Transgenic line | Gateway | Primers |
|---|---|---|
| Tg(Xla.Tubb2:QF2;he1.1:mCherry)$^{c663}$ | P5’E: Xla.Tubb2 | Forward: 5’-GGGGACAACTTTGTATAGAAAAGTTGTCTAGACCCTGTCTGTTCCTGA-3’<br>Reverse: 5’-GGGGACTGCTTTTTTGTACAAACTTGAAGCTTGATTGGGTTGAGTCTT-3’ |
| | pME: QF2-pA | Forward: 5’ -GGGGACAAGTTTGTACAAAAAAGCAGGCTCAACATGCCACCCAAGCG -3’<br>Reverse: 5’ -GGGGACCACTTTGTACAAGAAAGCTGGGTAAAAAACCTCCCACACCTCC -3’ |
| | P3’E: SV40pA;HE1.1:mCherry | Forward: 5’ -GGGGACAGCTTTCTTGTACAAAGTGGGATCCAGACATGATAAGATACAT -3’<br>Reverse: 5’ -GGGGACAACTTTGTATAATAAAGTTGCCATAGAGCCCACCGCAT -3’ |
| Tg(QUAS:mApple-CAAX; he1.1:mCherry)$^{c636}$ | P5’E: QUAS Through regular cloning | QUAS sequence: GGGTAATCGCTTATCCTCGGATAAACAATTATCCTCACGGGTAATCGCTTATCCGCTCGGGTAATCGCTTATCCTCGGGTAATCGCTTATCCTT<br>Carp promoter sequence: GCAAGGGTCGACTCTAGAGGGTATATAATGGATCCCATCGCGTCTCAGCCTCACTTTGAGCTCCTCCACACGAATTCCCTCGACCTCGAAGACGCGT |
| | pME: mApple-CAAX | Tol2kit v2.0 (#768) from Kristen Kwan |
| | P3’E: SV40pA;HE1.1:mCherry | Same as above |
| Tg(QUAS:GCaMP6f)$^{c587}$ | P5’E: QUAS | Same as above |
| | pME: GCaMP6f | Forward: 5’-GGGGACAAGTTTGTACAAAAAAGCAGGCTACGCCGCCACCATGGGTTCTCATCATCATCATCATCAT- 3’ |

| | | |
|---|---|---|
| | | Reverse: 5’-GGGGACCACTTTGTACAAGAAAGCTGGGTTCACTTCGCTGTCATCATTTGTAC-3’ |
| | P3’E: polyA | Tol2kit v1.2 (#302) from Chi-Bin Chien |
| Tg(QUAS:BoTxBLC-GFP)$^{c605}$ | P5’E: QUAS | Same as above |
| | pME: BoTxBLC-GFP | Forward: 5’-GGGGACAAGTTTGTACAAAAAAGCAGGCTATGCCCGTGACAATTAACAATT-3’<br>Reverse: 5’-GGGGACCACTTTGTACAAGAAAGCTGGGTTCACTTGTACAGCTCGTCCA-3’ |
| | P3’E: polyA | Tol2kit v1.2 (#302) from Chi-Bin Chien |
| Tg(QUAS:loxP-mCherry-loxP-GFP-CAAX)$^{c679}$ | P5’E: QUAS | Same as above |
| | pME: loxP-mCherry-stop-loxP | Tol2kit v2.0 (#759) from Kristen Kwan |
| | P3’E: GFP-CAAX-pA | Forward: 5’-GGGGACAGCTTTCTTGTACAAAGTGGAAATGGTGAGCAAGGGCGAG-3’<br>Reverse: 5’-GGGGACAACTTTGTATAATAAAGTTGAAAAAACCTCCCACACCTCCCCC-3’ |
| Tg(QUAS:loxP-mCherry-loxP-BoTxBLC-GFP)$^{c674}$ | P5’E: QUAS | Same as above |
| | pME: loxP-mCherry-stop-loxP | Tol2kit v2.0 (#759) from Kristen Kwan |
| | P3’E: BoTxBLC-GFP-stop | Forward: 5’-GGGGACAGCTTTCTTGTACAAAGTGGAAATGCCCGTGACAATTAACAATT-3’<br>Reverse: 5’-GGGGACAACTTTGTATAATAAAGTTGTCACTTGTACAGCTCGTCCA-3’ |

**Key resources table**

| Reagent type (species) or resource | Designation | Source or reference | Identifiers | Additional information |
|---|---|---|---|---|
| Genetic reagent (*Danio rerio*) | Tg(lratd2a:QF2)$^{c601}$ | This paper | | Transgenic |
| Genetic reagent (*Danio rerio*) | Tg(slc5a7a:Cre)$^{c662}$ | This paper | | Transgenic |
| Genetic reagent (*Danio rerio*) | Tg(Xla.Tubb2:QF2;he1.1: mCherry)$^{c663}$ | This paper | | Transgenic |
| Genetic reagent (*Danio rerio*) | *Tg(QUAS:GCaMP6f)*$^{c587}$ | This paper | | Transgenic |
| Genetic reagent (*Danio rerio*) | *Tg(QUAS:mApple-CAAX;he1.1:mCherry)*$^{c636}$ | This paper | | Transgenic |
| Genetic reagent (*Danio rerio*) | *Tg(QUAS:loxP-mCherry-loxP-GFP-CAAX)*$^{c679}$ | This paper | | Transgenic |
| Genetic reagent (*Danio rerio*) | *Tg(QUAS:loxP-mCherry-loxP-BoTxBLC-GFP)*$^{c674}$ | This paper | | Transgenic |
| Genetic reagent (*Danio rerio*) | *Tg(-10lhx2a:gap-EYFP)*$^{zf177}$ | Miyasaka et al., 2009 | RRID:ZFIN_ZDB-GENO-100504-13 | Transgenic |
| Genetic reagent (*Danio rerio*) | *tcf7l2*$^{zf55}$ | Muncan et al., 2007 | RRID:ZFIN_ZDB-GENO-071217-3 | Mutant |
| Genetic reagent (*Danio rerio*) | *bsx*$^{m1376}$ | Schredelseker el al., 2018 | RRID:ZFIN_ZDB-GENO-180802-1 | Mutant |
| Chemical compound, drug | alpha-Bungarotoxin | Invitrogen | Cat# B-1601 | 1 mg/ml |
| Chemical compound, drug | Cadaverine | Sigma-Aldrich | Cat# 33211 | 100 µM |
| Chemical compound, drug | Chondroitin sulfate sodium salt from shark cartilage | Sigma-Aldrich | Cat# C4384 | 100 µg/ml |
| Chemical compound, drug | T7 Endonuclease I | NEB | Cat# M0302L | |
| Chemical compound, drug | MAXIscript™ T7 Transcription Kit | Invitrogen | Cat# AM1312 | |
| Chemical compound, drug | mMASSAGE mMACHINE T3 Transcription Kit | Invitrogen | Cat# AM1348 | |
| Chemical compound, drug | DIG RNA Labeling Mix | Roche | Cat# 11277073910 | |
| Chemical compound, drug | 5-bromo-4-chloro-3-indolyl-phosphate, 4-toluidine salt (BCIP) | Roche | Cat# 11383221001 | |
| Chemical compound, drug | 4-Nitro blue tetrazolium chloride, solution (NBT) | Roche | Cat# 11383213001 | |

| | | | | |
|---|---|---|---|---|
| Chemical compound, drug | 2-(4-Iodophenyl)-3-(4-nitrophenyl)-5-phenyltetrazolium Chloride (INT) | FisherScientific | Cat# I00671G | |
| Antibody | Anti-Digoxigenin-AP, Fab fragments antibody | Roche | Cat# 11093274910, | 1:5000 |
| Antibody | Anti- Fluorescein -AP, Fab fragments antibody | Roche | Cat# 11426338910 | 1:5000 |
| Recombinant DNA reagent | Plasmid: Gbait-hs-Gal4 | Kimura et al., 2014 | N/A | |
| Recombinant DNA reagent | Plasmid: Gbait-hsp70-QF2-SV40pA | This paper | Addgene Plasmid #122563 | |
| Recombinant DNA reagent | Plasmid: Gbait-hsp70-Cre-SV40pA | This paper | Addgene Plasmid #122562 | |
| Recombinant DNA reagent | Plasmid: pDR274 | Hwang et al., 2013 | Addgene Plasmid #42250 | |
| Recombinant DNA reagent | Plasmid: GFP sgRNA | Auer et al., 2014 | N/A | |
| Recombinant DNA reagent | Plasmid: pT3TS-nCas9n | Jao et al., 2013 | #46757 | |
| Recombinant DNA reagent | pGEM®-T Easy Vector | Promega | Catalog# A1360 | |
| Software, Algorithm | Fiji | Schindelin et al., 2012 | | https://imagej.net/Fiji |
| Software, Algorithm | MATLAB | The MathWorks | | https://www.mathworks.com/ |
| Software, Algorithm | Excel | Microsoft | | http://products.office.com/en-us/excel |
| Software, Algorithm | ZebraLab | Viewpoint Life Sciences | | http://www.viewpoint.fr/en/home |
